# Kolmogorov-Arnold Networks for Genomic Tasks

**DOI:** 10.1101/2024.12.08.627375

**Authors:** Oleksandr Cherednichenko, Maria Poptsova

## Abstract

Kolmogorov-Arnold Networks (KANs) emerged as a promising alternative for multilayer perceptrons in dense fully connected networks. Multiple attempts have been made to integrate KANs into various deep learning architectures in the domains of computer vision and natural language processing. Integrating KANs into deep learning models for genomic tasks has not been explored. Here, we tested linear KANs (LKANs) and convolutional KANs (CKANs) as replacement for MLP in baseline deep learning architectures for classification and generation of genomic sequences. We used three genomic benchmark datasets: Genomic Benchmarks, Genome Understanding Evaluation, and Flipon Benchmark. We demonstrated that LKANs outperformed both baseline and CKANs on almost all datasets. CKANs can achieve comparable results but struggle with scaling over large number of parameters. Ablation analysis demonstrated that the number of KAN layers correlates with the model performance. Overall, linear KANs show promising results in improving the performance of deep learning models with relatively small number of parameters. Unleashing KAN potential in different SOTA deep learning architectures currently used in genomics requires further research.

## 1. Introduction

Deep learning models have been successfully applied to a wide range of genomic tasks including prediction of variant effect (Zhou & Troyanskaya, 2015),(Avsec et al., 2021), and location of functional genomic elements, such as promoters (Oubounyt et al., 2019a), enhancers (Le et al., 2021), transcription factor binding sites (Quang & Xie, 2019), histone marks (Yin et al., 2021), (Wen et al., 2024), flipons (Beknazarov et al., 2020), (Umerenkov et al., 2023), splice sites (de Sainte Agathe et al., 2023), and others. Neural networks also proved to be efficient in the field of RNA biology (see (Sato & Hamada, 2023) and (Hwang et al., 2024) for review) predicting RNA-protein binding (Shen et al., 2020), RNA structure (Franke et al., 2024), non-coding RNA (Georgakilas et al., 2020).

The evolution of deep learning applications in genomics followed the path of the evolution of deep learning architectures. It started with pioneering application of Convolutional Neural Networks (CNNs) (Zhou & Troyanskaya, 2015; Chen et al., 2022) and Reccurent Neural Networks (RNNs) to DNA sequences (Shen et al., 2018), and then continued with Transformer-based Large Language Models (LLMs) such as DNABERT (Ji et al., 2021), DNABERT2 (Zhou et al., 2024b), Nucleotide Transformer (Dalla-Torre et al., 2023). DNABERT was trained on one reference human genome while DNABERT2 and Nucleotide transformer were trained on the multi-species genome comprising up to 40B of bases pairs. After training foundation models are fine-tuned on different downstream tasks however even with parameter efficient fine-tuning techniques, resource capability can impose significant limitations on the task realization.

Transformers require substantial computational resources due to quadratic attention scaling, and the next architecture Hyena, based on long convolutions, incorporated longer context up to 1M nucleotides by scaling subquadratically, which resulted in HyenaDNA (Nguyen et al., 2023). Another promising alternative to overcome transformers’ computational inefficiency is Mamba architecture (Gu & Dao, 2023) which lie at the basis of Caduceus model in genomics (Schiff et al., 2024). Both Hyena and Mamba achieved a certain tradeoff between computational resources and performance quality. In comparison to the latest genomic foundational model DNABERT2, HyenaDNA has 10x less parameters and on some genomic tasks outperforms DNABERT2.

With ever-growing size of deep learning architectures, it has been a challenge to find smaller architectures that would perform equally well. Here we aim to investigate the potential of small neural networks that utilize only few MLPs layers with recent innovations in deep learning such as Kolmogorov-Arnold Networks (KANs) (Liu et al., 2024b).

The neural networks discussed above have one thing in common: they rely on the universal approximation theorem. One of the recent advances in deep learning architectures, Kolmogorov-Arnold Network (Liu et al., 2024b), leverages the Kolmogorov-Arnold theorem to incorporate splines into a neural network architecture, offering a compelling alternative to traditional Multi-Layer Perceptrons (MLPs). KANs have already been successfully applied in different areas such as mechanics (Liu et al., 2024a; Abueidda et al., 2024), computer vision (Bodner et al., 2024; Drokin, 2024; Yang & Wang, 2024; Aghaei, 2024; Cheon, 2024), NLP (Yu et al., 2024; Genet & Inzirillo, 2024b), time series (Genet & Inzirillo, 2024b;a; Zhou et al., 2024a), physics (Liu et al., 2024b;a; Ta, 2024), speech enhancement (Xu et al., 2024a), molecular representations (Li et al., 2024; Nagai & Okumura, 2024), and recommendation systems (Park et al., 2024). Inspired by this advancement, multiple modifications have emerged that attempt to overcome various issues associated with the spline-based approach, namely computational overhead and a large number of trainable parameters.

In the presented study we aim at evaluating the potential of KAN-based models in genomics. Given the limitations in computational resources, we applied KAN to simple convolutions and dense networks with a relatively small number of parameters. We tested KANs to the task of classification and generation by replacing layers in the baseline model with Linear KAN layers (LKAN) and Convolutional KAN layers (CKAN). The test was performed on three benchmark datasets: Genomic Benchmarks (Gresova et al., 2023), Genome Understanding Evaluation (GUE) and our collection of datasets for flipons. or non-B DNA structures (Herbert, 2024).

### 1.1. Kolmogorov-Arnold Networks

#### 1.1.1. Linear Kolmogorov-Arnold Networks

Here we briefly introduce KAN and present key features of this architecture. Multi-Layer Perceptrons (Hornik et al., 1989) are inspired by the universal approximation theorem that states that a feed-forward network with a single hidden layer containing a finite number of neurons can approximate continuous functions on compact subsets of R^*d*^. Kolmogorov-Arnold Network (Liu et al., 2024b) focuses on the Kolmogorov-Arnold representation theorem (Kolmogorov, 1956) which states that any multivariate continuous function can be represented as a composition of univariate functions and the addition operation

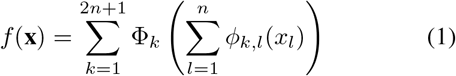

where *ϕ*_*k,l*_ are univariate functions that map each input variable *x*_*l*_ as follows: *ϕ*_*k,l*_ : [0, 1] *→* ℝ. The authors of the original KAN paper proposed how to extend the KAN application to deep learning networks by stacking KAN layers.

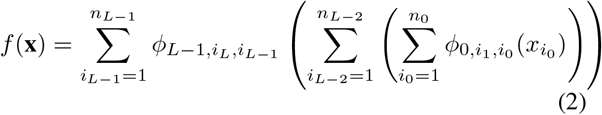

Similarly it is possible to rewrite it in a way

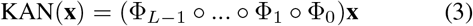

Implementation of Kolmogorov-Arnold Networks is provided by the authors of (Liu et al., 2024b), however there are several aspects that could limit the scalability of KANs: a large number of trainable parameters, training time, and inference time. On top of that, in (Yu et al., 2024) the authors claim that weights have different initialization that makes it impossible to keep variance-preserving initialization (Yang & Wang, 2024). To overcome these issues, in (Yang & Wang, 2024) the authors suggest an efficient implementation of KANs with the straightforward matrix multiplication and L1 regularization for the model’s weights.

#### 1.1.2. Kolmogorov-Arnold Convolutions

Kolmogorov-Arnold Convolutions are proposed in (Bodner et al., 2024) and designed to be similar to CNNs. The difference is that the convolutional layers are replaced by KAN convolutional layers, and, after flattening, one can use either KAN or MLP. The advantage of the Convolutional KANs is that they have significantly fewer parameters compared to other architectures. This is provided by the network architecture, because B-Splines are capable of smooth representations of aribtrary activation functions, which cannot be determined with ReLU. The authors concluded that Kolmogorov-Arnold Convolutions are capable of achieving high performance as compared to original convolutional networks (Drokin, 2024).

### 1.2. Genomics Tasks

#### 1.2.1. DNA classification

Classification of DNA sequences is one of the most common tasks in genomics solved with deep learning models. In this work, we perform classification on three large benchmark datasets: Genomic Benchmarks (Gresova et al., 2023), Genomic Understanding Evaluation (Zhou et al., 2024b), and Flipon Benchmark assembled by us. For each of the datasets there available a train and a test sets.

#### 1.2.2. DNA generation

Another important application of deep learning in genomics is generative modeling. Generative models are usually used in data augmentation approaches to generate new samples from the learned distribution.

There are many types of generative models, but here we focus on the most popular models for DNA sequence generation: denoising diffusion implicit model (DDIM) (Song et al., 2021) and generative adversarial network (GAN) (Arjovsky et al., 2017). In generative approaches, first, a generative model is trained on a train set, and the trained model produces synthetic samples of the same size as the test set. Secondly, synthetic data is supplied to a classifier trained on the real data to evaluate how good are generative models in capturing main patterns of the real data.

We will use the Flipons collection (Kouzine, 2017; Shin, 2016; Qian, 2024; Zhang et al., 2019; Chambers et al., 2015; Lyu et al., 2016) for this task (see Methods).

## 2. Methods

### 2.1. Models

We use pytorch efficient implementation of KAN (EKAN) (Yang & Wang, 2024) as linear KAN layers. We use KAN-Conv (Bodner et al., 2024) implementation as KAN convolutional neural network. We incorporate KAN layers into convolutional neural network that takes its core from LegNet (Penzar et al., 2023). We compare the impact of KAN layers by replacing MLP modules with KANs (see Tables 1–5). To make a comparison more fair and to exclude model size effect, in all the experiments for classification we used models of equal size: 22M parameters. We used the following notation for our tested modesl: Baseline for LegNet-based convolutional neural network, LKAN for LegNet-based convolutional neural network (Tan & Le, 2024) with incorporated Linear KANs, and CKAN for LegNet-based convolutional neural network with incorporated Convolutional KANs. Since CKANs scales with larger grid size (see Results), combining CKAN and LKAN in our architecture was impossible due to limitations in resources, thus hybrid CKAN-LKAN architectures were not conducted.

**Table 1.** KAN performance on classification task for Genomic Benchmarks based on MCC-score averaged over 5-folds. Best values are in **bold**, second best value is in *italics*.

| DATA SET | BASELINE | LKAN | CKAN |
| --- | --- | --- | --- |
| DMEE | 70.2± 0.5 | **73.2± 0.2** | 64.8± 1.3 |
| DCIS | 81.1± 0.5 | **85.5± 1.4** | 81.7± 1.2 |
| DHW | 89.2± 0.5 | **90.2± 0.5** | 88.1± 0.8 |
| DES | **27.3± 0.9** | 28.9± 0.6 | 25.3± 0.6 |
| HEC | 66.6± 0.8 | **68.2± 0.5** | 67.9± 0.8 |
| HEE | 63.2± 0.5 | **69.9± 0.5** | 59.0± 1.0 |
| HER | **90.0± 0.8** | 89.3± 0.8 | 88.4± 0.6 |
| HNP | 87.9± 0.9 | **88.8± 0.9** | 85.0± 1.0 |
| HOE | 70.8± 0.6 | **74.4± 0.7** | 68.9± 1.0 |

In experiments with DNA sequence generation we utilized denoising diffusion implicit model (Song et al., 2021) designed specifically for DNA (DaSilva, 2024). We replaced linear layers with KAN layers in Unet like architecture.

### 2.2. Datasets

#### 2.2.1. Genomic Benchmarks

We use the following notations for all datasets: DMEE - dummy mouse enhancers ensemble, DCIS - demo coding vs intergenomic seqs, DHW - demo human or worm, DES - drosophila enhancers stark, HEC - human enhancers cohn, HEE - human enhancers ensembl, HER - human ensembl regulatory, HNP - human nontata promoters, HOE - human ocr ensembl.

#### 2.2.2. Flipons, Or NON-B DNA STRUCTURES

Inspired by Genomic Benchmarks we created a collection of datasets for detecting flipons (Herbert, 2024), or non-B DNA structures: Z-flipons, or Z-DNA (Kouzine, 2017; Shin, 2016); G-flipons, or G-quadruplexes (GQs) (Qian, 2024; Zhang et al., 2019; Chambers et al., 2015; Lyu et al., 2016); H-flipons, or H-DNA (triplexes) (Kouzine, 2017). We provide train and test sets trying to prevent data leakage. Full description of datasets are available in Supplementary Table 1. Similar to Genomic Benchmarks datasets we use the following notations for all flipons datasets: ENDO - Endoquad GQs; G4seq - G4-seq experimental dataset for GQs; G4ChIP - G4 ChIP-seq experimental data set for GQs; G4cut - G4 CUT&Tag experimental dataset for GQs; ZKOU Kouzine et al. experimental dataset for Z-DNA; ZShin - Shin et al. ChIP-seq experimental dataset for Z-DNA; Hkou Kouzine experimental dataset for H-DNA.

#### 2.2.3. Genome Understanding Evaluation

We follow (Zhou et al., 2024b) authors in evaluation on Genome Understanding Evaluation (GUE) and extended version (GUE+) to study models’ performance on a wider range of genomic tasks: prediction of promoter, core promoter, splice sites, epigenetic marks, transcription factor binding sites on human and mouse, and covid variant classification.

## 3. Results

### 3.1. Benchmarks on DNA classification

#### 3.1.1. Genomic Benchmarks

KAN performance in comparison to the baseline model for nine datasets from Genomic Benchmarks (Gresova et al., 2023) are presented in Table 1 (MCC score averaged from 5 folds). Baseline CNN model is of equal size with LKAN and CKAN. Other classification metrics such as accuracy, ROC-AUC, precision, recall, and F1 are provided in Supplementary Tables 2-14. We observe that both LKAN and CCKAN improve the quality of the model, yet CKAN requires higher value of grid size parameter.

#### 3.1.2. Flipons, or non-b dna structures

KAN performance on seven datasets from Flipon collection (Kouzine, 2017; Shin, 2016; Qian, 2024; Zhang et al., 2019; Chambers et al., 2015; Lyu et al., 2016) is presented in Table 2. Baseline CNN model is of equal size with LKAN and CKAN.

**Table 2.**
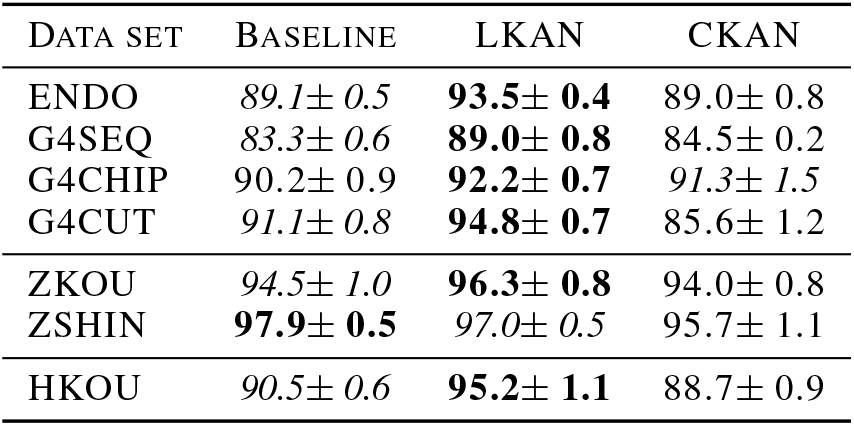
KAN performance on classification task for flipons based on MCC-score averaged over 5-folds. Best values are in **bold**, second best value is in *italics*.

#### 3.1.3. Genome Understanding Evaluation

Results of KAN performance in comparison to the baseline model for nine datasets from Genome Understanding Evaluation (Zhou et al., 2024b) are presented in Table 3 (MCC score averaged from 5 folds). Baseline CNN model is of equal size with LKAN and CKAN.

**Table 3.** KAN performance on classification task for GUE based on MCC averaged over 5-folds. Best values are in **bold**, second best value is in *italics*.

| DATA SET | BASILINE | LKAN | CKAN |
| --- | --- | --- | --- |
| H3 | 70.7 ± 1.5 | **71.3 ± 1.0** | 69.4 ± 1.5 |
| H3K14AC | 40.2 ± 0.8 | **43.2 ± 0.7** | 41.0 ± 1.0 |
| H3K36ME3 | 44.3 ± 0.9 | **47.7 ± 0.6** | 42.9 ± 0.9 |
| H3K4ME1 | 35.6 ± 0.4 | **40.0 ± 0.8** | 35.8 ± 1.1 |
| H3K4ME2 | 27.2 ± 0.5 | **29.3 ± 0.5** | 24.3 ± 1.2 |
| H3K4ME3 | 28.4 ± 0.5 | **29.8 ± 0.4** | 27.2 ± 0.7 |
| H3K79ME3 | 60.8 ± 0.9 | **62.1 ± 0.8** | 61.1 ± 1.5 |
| H3K9AC | 50.5 ± 0.6 | **51.9 ± 0.5** | 50.0 ± 0.9 |
| H4 | 75.6 ± 0.9 | **77.2 ± 0.7** | 70.9 ± 0.8 |
| H4AC | 33.9 ± 0.8 | **38.1 ± 0.8** | 36.2 ± 0.9 |
| PD ALL | 75.9 ± 1.1 | **81.2 ± 1.4** | 77.2 ± 1.2 |
| PD NOTATA | 88.2 ± 1.1 | **89.9 ± 0.7** | 87.1 ± 1.2 |
| PD TATA | 56.1 ± 0.5 | **58.2 ± 0.8** | 57.9 ± 0.9 |
| TF HUMAN 1 | 69.9 ± 1.0 | **72.4 ± 0.8** | 66.3 ± 1.3 |
| TF HUMAN 2 | 49.6 ± 0.4 | **54.3 ± 0.7** | 47.2 ± 0.7 |
| TF HUMAN 3 | 43.2 ± 0.5 | **46.8 ± 0.5** | 44.4 ± 0.9 |
| TF HUMAN 4 | 72.6 ± 0.8 | **72.9 ± 0.5** | 71.3 ± 0.9 |
| TF HUMAN 0 | 65.1 ± 0.7 | **67.4 ± 0.6** | 65.5 ± 0.9 |
| CPD ALL | 64.8 ± 0.9 | **68.2 ± 0.6** | 66.2 ± 1.1 |
| CPD NOTATA | 64.5 ± 0.7 | **67.2 ± 0.8** | 63.1 ± 1.1 |
| CPD TATA | 72.3 ± 0.9 | **78.2 ± 0.9** | 74.7 ± 1.2 |
| TF MOUSE 1 | 81.0 ± 0.7 | **82.8 ± 0.6** | 80.1 ± 0.8 |
| TF MOUSE 2 | 79.8 ± 0.6 | **85.5 ± 0.8** | 81.2 ± 0.7 |
| TF MOUSE 3 | 77.6 ± 1.2 | **81.4 ± 0.7** | 79.2 ± 0.9 |
| TF MOUSE 4 | 42.6 ± 0.7 | **48.7 ± 0.5** | 40.1 ± 0.7 |
| TF MOUSE 0 | 38.2 ± 0.7 | **40.1 ± 0.6** | 39.0 ± 0.9 |
| VIRUS | 50.3 ± 0.9 | **58.2 ± 1.5** | 50.5 ± 0.6 |
| SPICE | 78.2 ± 0.8 | **80.0 ± 0.8** | 75.2 ± 0.7 |

**Table 4.** KAN performance on classification task for GUE+ based on MCC averaged over 5-folds. Best values are in **bold**, second best value is in *italics*.

| DATA SET | BASILINE | LKAN | CKAN |
| --- | --- | --- | --- |
| GM12878 | 36.6 ± 1.1 | **41.2 ± 0.9** | 37.0 ± 1.2 |
| HELA-S3 | 34.3 ± 1.0 | **40.8 ± 1.2** | 31.3 ± 0.9 |
| HUVEC | 37.2 ± 0.7 | **39.6 ± 0.9** | 30.2 ± 0.8 |
| IMR90 | 34.5 ± 0.9 | **42.0 ± 0.9** | 33.3 ± 1.3 |
| K692 | 38.2 ± 0.9 | **44.4 ± 0.7** | 37.9 ± 1.1 |
| NHEK | 33.7 ± 1.1 | **40.1 ± 0.9** | 34.4 ± 0.6 |
| FUNGI | 51.1 ± 0.6 | **55.5 ± 1.0** | 50.4 ± 1.2 |
| VIRUS | 20.7 ± 0.5 | **35.3 ± 1.9** | 22.1 ± 1.8 |

Considering results for the classification tasks we observe that Linear KAN scales better than Convolutional KAN of the same size. We present visualization of averaged model’s performance for models of various size in Figure 3. Additionally, we provide results of experiments on how grid size parameter affects the performance of LKAN and the number of total parameters along with the time per batch training. We evaluated these results on GUE datasets.

**Figure 1.**
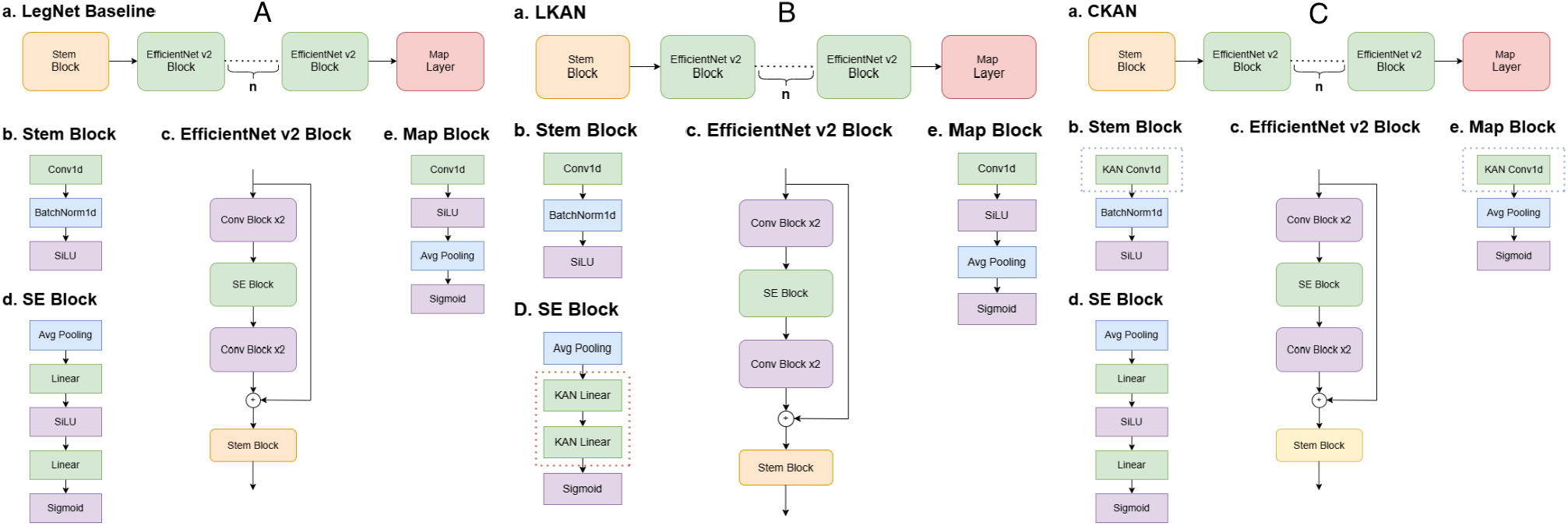
**A. LegNet-based Baseline. a**. Architecture of Baseline LegNet-based convolutional neural network. The Number of EfficientNet-like block are defined as *n* and we use *n* = 6. Other values were tested in Ablation study. **b**. Stem block incorporated in Baseline architecture. **c**. Main convolutional EfficientNet-like (Tan & Le, 2024) block incorporated in Baseline architecture. Plus sign refers to per channel concatenation. **d**. Mid SE block with bilinear layer incorporated in Baseline architecture. **e**. Final block incorporated in Baseline architecture to access final prediction. **B. LegNet-based LKAN. a**. Architecture of Baseline LegNet-based convolutional neural network with LKAN. The number of EfficientNet like block are defined as *n* and we use *n* = 6. Other values were tested in Ablation study. **b**. Stem block incorporated in Baseline architecture. **c**. Main convolutional EfficientNet-like (Tan & Le, 2024) block incorporated in Baseline architecture. Plus sign refers to per channel concatenation. **d**. Mid SE block with replaced LKAN layers incorporated in Baseline architecture. **e**. Final block incorporated in Baseline architecture to access a final prediction. **C. LegNet-based CKAN. a**. Architecture of Baseline LegNet-based convolutional neural network with CKAN. The number of EfficientNet-like block are defined as *n* and we use *n* = 6. Other values were tested in Ablation study. **b**. Stem block with replaced CKAN layer incorporated in Baseline architecture. **c**. Main convolutional EfficientNet-like (Tan & Le, 2024) block incorporated in Baseline architecture. Plus sign refers to per channel concatenation. **d**. Mid SE block with bilinear layer incorporated in Baseline architecture. **e**. Final block with replaced CKAN layer incorporated in Baseline architecture to access a final prediction.

**Figure 2.**
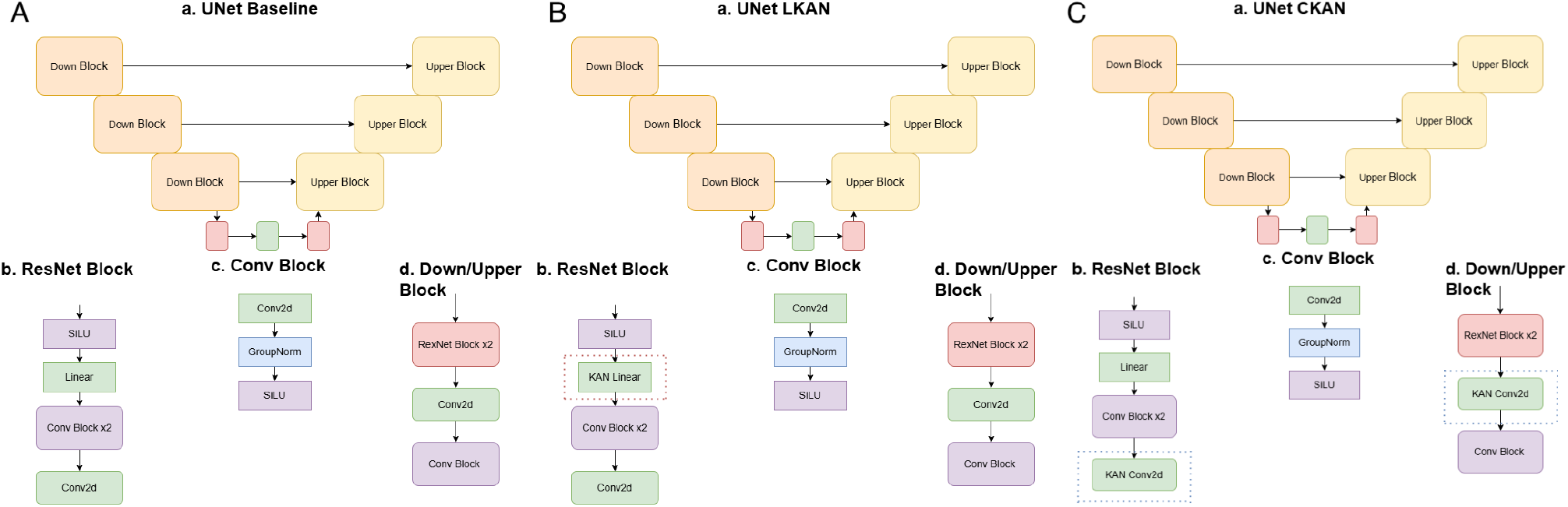
**A. Unet-based Baseline a**. Architecture of Baseline DNA-Diffusion Unet-based model. Model’s core consists of upper and down blocks. **b**. Base ResNet block gathered from Convolutional Blocks. **c**. Convolutional Block is incorporated into base ResNet block and at the end of Upper and Down blocks. **d**. Structure of Down and Upper Blocks with ResNet Blocks. **B. Unet-based LKAN a**. Architecture of Linear Kolmogorov Arnold Network (LKAN) modification of DNA-Diffusion Unet-based model. Model’s core consists of upper and down blocks. **b**. Base ResNet block gathered from Convolutional Blocks. Dense layer in midst replaced with Linear KAN. **c**. Convolutional Block is incorporated into base ResNet block and at the end of Upper and Down blocks. **d**. Structure of Down and Upper Blocks with ResNet Blocks. **C. Unet-based CKAN a**. Architecture of Convolutional Kolmogorov Arnold Network (CKAN) modification of DNA-Diffusion Unet-based model. Model’s core consists of upper and down blocks. **b**. Base ResNet block gathered from Convolutional Blocks with replaced CKANs. **c**. Convolutional Block is incorporated into base ResNet block and at the end of Upper and Down blocks. **d**. Structure of Down and Upper Blocks with ResNet Blocks. Convolutional layer replaced with CKAN.

**Figure 3.**
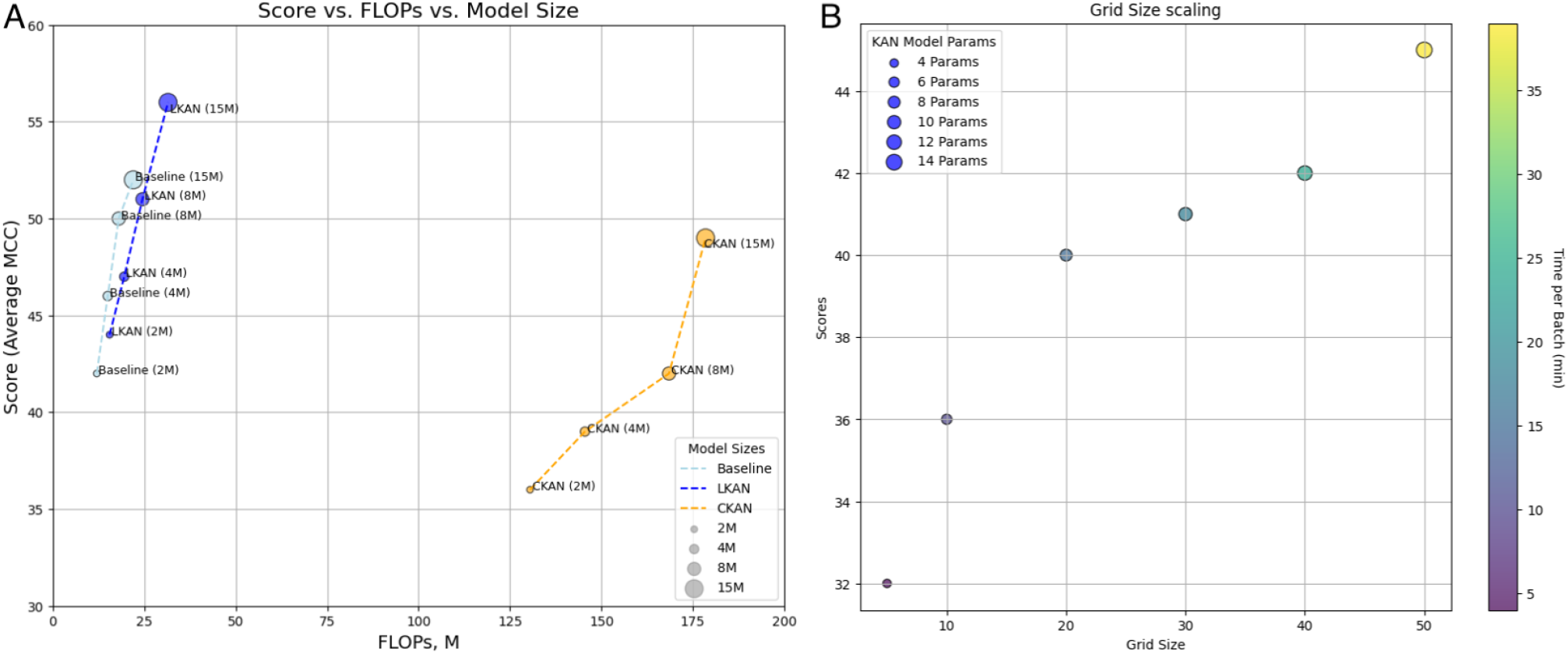
Scaling of models. **A**. Comparison of the tested models of various sizes, evaluated on different collections of datasets. Scores are calculated as average MCC and FLOPs are calculated per input batch. We observe that Linear KAN performs the best and scales the same as Baseline. In contrast, CKAN requires more computations per second and shows relatively poor performance. Achieving better performance with CKAN requires heavier model in terms of parameters. **B**. Linear KAN grid size scaling on model’s parameters and training time. We used GUE+ collection for this experiment. As one can see, grid size significantly increases training time.

### 3.2. Benchmarks on DNA generative models

To test KAN’s capability in producing synthetic DNA sequences we incorporated Linear KAN layers into denoising Unet from DNA Diffusion framework (DaSilva, 2024).

We compared final loss for various models (Table 5) and observed that LKAN is capable of reaching the lowest validation loss given equal amount of parameters with baseline Unet. CKAN requires x2 parameters to achieve the comparable metrics.

**Table 5.** KANs for DNA generation: KAN loss (SP-MSE) comparison on flipon generation. Best values are in **bold**, second best value is in *italics*

| TASK | UNET (410M) | LKAN (440M) | CKAN (820M) |
| --- | --- | --- | --- |
| G4 | 0.0334 | **0.0276** | 0.0340 |
| ZDNA | 0.0301 | **0.0222** | 0.0299 |
| HDNA | 0.0388 | **0.0281** | 0.0379 |

Additionally, we observe that KAN affects training stability of Unet, as it is presented in Figure 4. Both trainings utilize exponential moving average for smoother performance. We conclude that Linear KANs provide better performance than Convolutional KANs, yet Convolutinal KANs requires higher values for grid size, which significantly heavies up the model in terms of parameters. For instance, CKAN requires 820M parameters while LKAN requires 440M to outperform baseline Unet.

We compare the performance of KAN in generative design by evaluating Kullback–Leibler divergence and Wasserstein distance. Since diffusion model require substantial resources for training and inference (see Supplementary Materials for details), we conduct these experiments only for flipon’s datasets (Cherednichenko & Poptsova, 2025). We compare the diversity calculated on edit distance within a sample of synthetic sequences for various models (Table 6). Higher values ensure more diversity within a sample. Trained generative models have captured the prior distributions of flipons based on calculated distances between real and generated distributions (Figures 3–4).

**Table 6.**
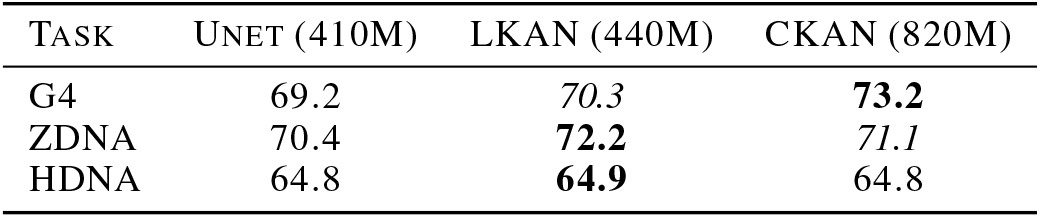
KANs for DNA generation: metrics of diversity for flipons. Best values are in **bold**, second best value is in *italics*.

**Figure 4.**
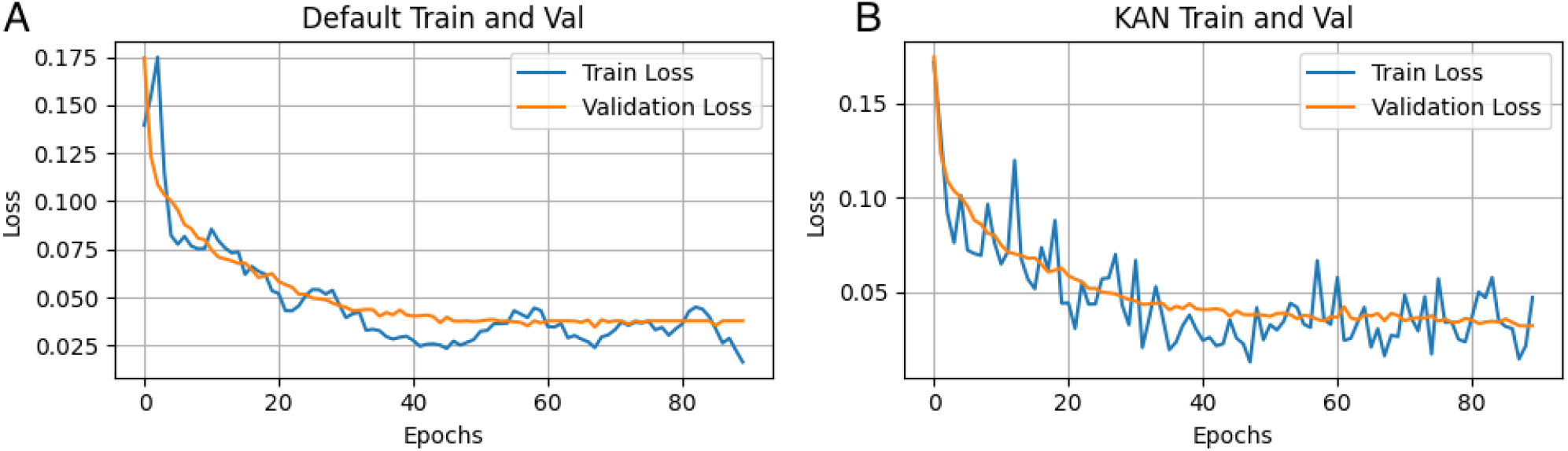
Loss functions of generative models. **A**. Baseline Unet with Exponential Moving Average (EMA). **B**. Unet with replaced MLP with Linear KAN (LKAN) also with EMA. While LKAN generally achieves better loss values, it is challenging to stabilize training, even with EMA it gets more spiky.

### 3.3. Ablation study

Additionally, we investigated the impact of the total amount of replaced MLP layers with KANs. For this experiment we used human enhancers cohn dataset from Genomic Benchmarks collection. Taking into consideration the total number of EfficientNet like blocks (Supplementary Table 15) we decided to determine if the same quality can be achieved by replacing only few blocks with LKANs. Model performance based on F1 score with the number of replaced MLP layers and the grid size is presented in Supplementary Table 15. We observe a steady increase in performance with the number of replaced layers. We also see that increasing grid size positively impacts the overall performance. Thus, we found that the best performance is achieved when all blocks are replaced with LKANs, however, with higher grid size values it is possible to achieve fair performance with *N* = 4 or *N* = 5 (see Supplementary Tables 15-16). Yet, increasing grid size makes model heavier, therefore the training process takes longer. Moreover, higher grid size is prone to certain overfit, when training metrics are close to high values while testing metrics remain constant (Supplementary Figure 1).

## 4. Discussions and Conclusions

In the presented study, we evaluated performance of the recently developed Kolmogorov-Arnold Networks in the domain of genomics. For that we used a wide range of datasets to cover different types of genomic functional elements including promoters, enhancers, histone marks, and others collected in two published genomic benchmark datasets and one novel benchmark dataset for flipons, or non-B DNA structures, assembled by us. For our baseline models i.e. models without KAN, we utilized LegNet-based convolutional network for DNA classification (Penzar et al., 2023) and denoising diffusion implicit model DNA-Diffusion with a core Unet module for generating synthetic sequences (DaSilva, 2024). MLP and convolutional layers in these baselines were replaced with linear and convolution KAN layers correspondingly.

Our benchmarking show that Linear KANs show promising results in replacing MLPs in a broad range of genomic tasks, while Convolutional KANs struggle with scaling over large number of parameters, which require more computational resources. In sequence generation we proved that all models, baseline and KANs, can learn distributions of the input data (Figures 5–6), however LKAN outperforms baseline and CKAN in terms of Loss, while CKAN requires x2 more parameters to achieve the same performance as LKAN. In diversity of generated samples both LKANs and CKANs outperform baseline.

**Figure 5.**
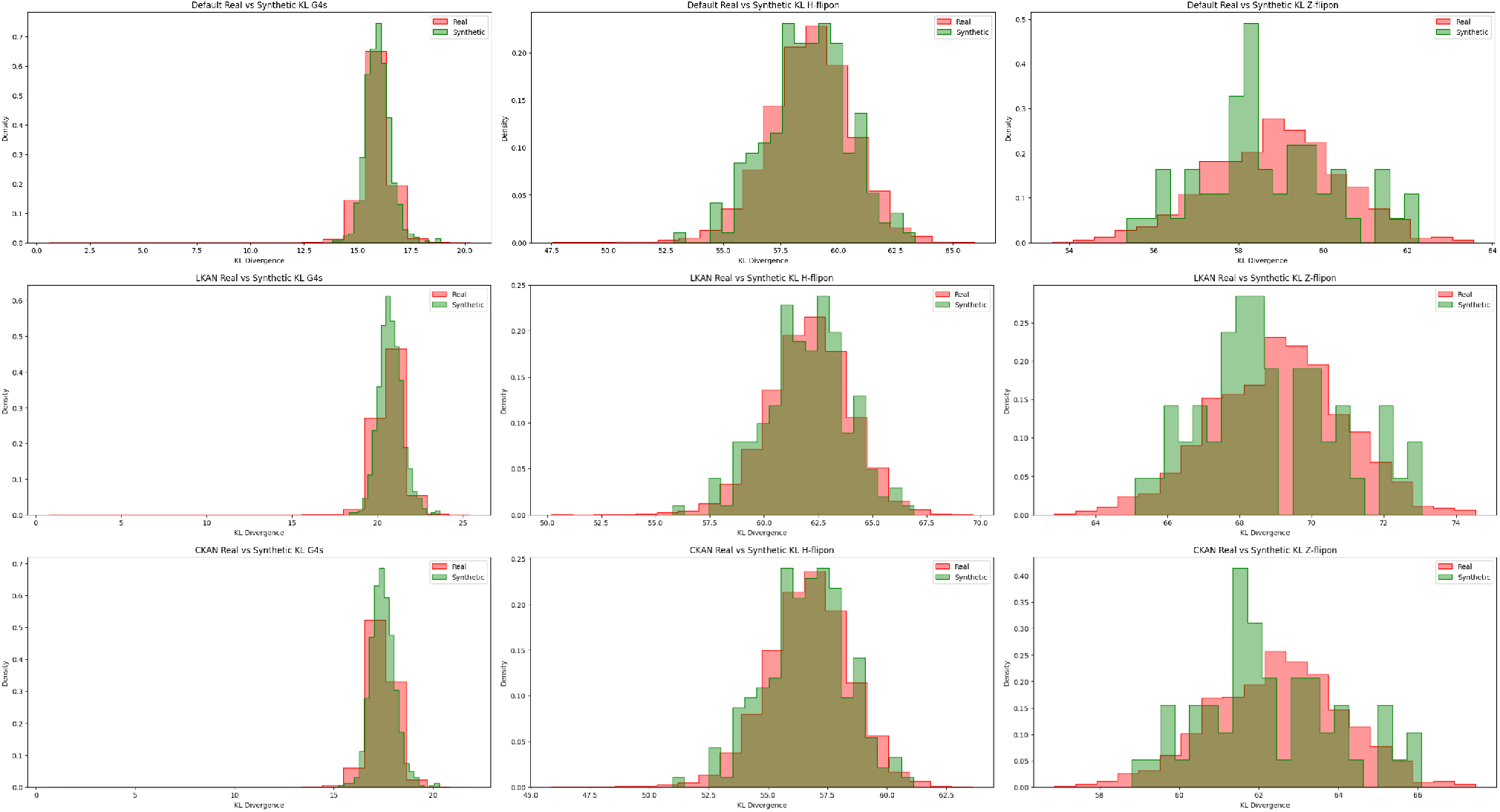
Distributions of Kullback–Leibler divergence (KL) for real and synthetic data. Synthetic data are produced by Baseline Unet (Default) and Unet with Linear KAN (LKAN). We conducted this experiment for flipon benchmark datasets and presnet distributions for Z-DNA, H-DNA and G4s. Results show that both generative models learned to approximate prior distribution as green and red distributions are close.

**Figure 6.**
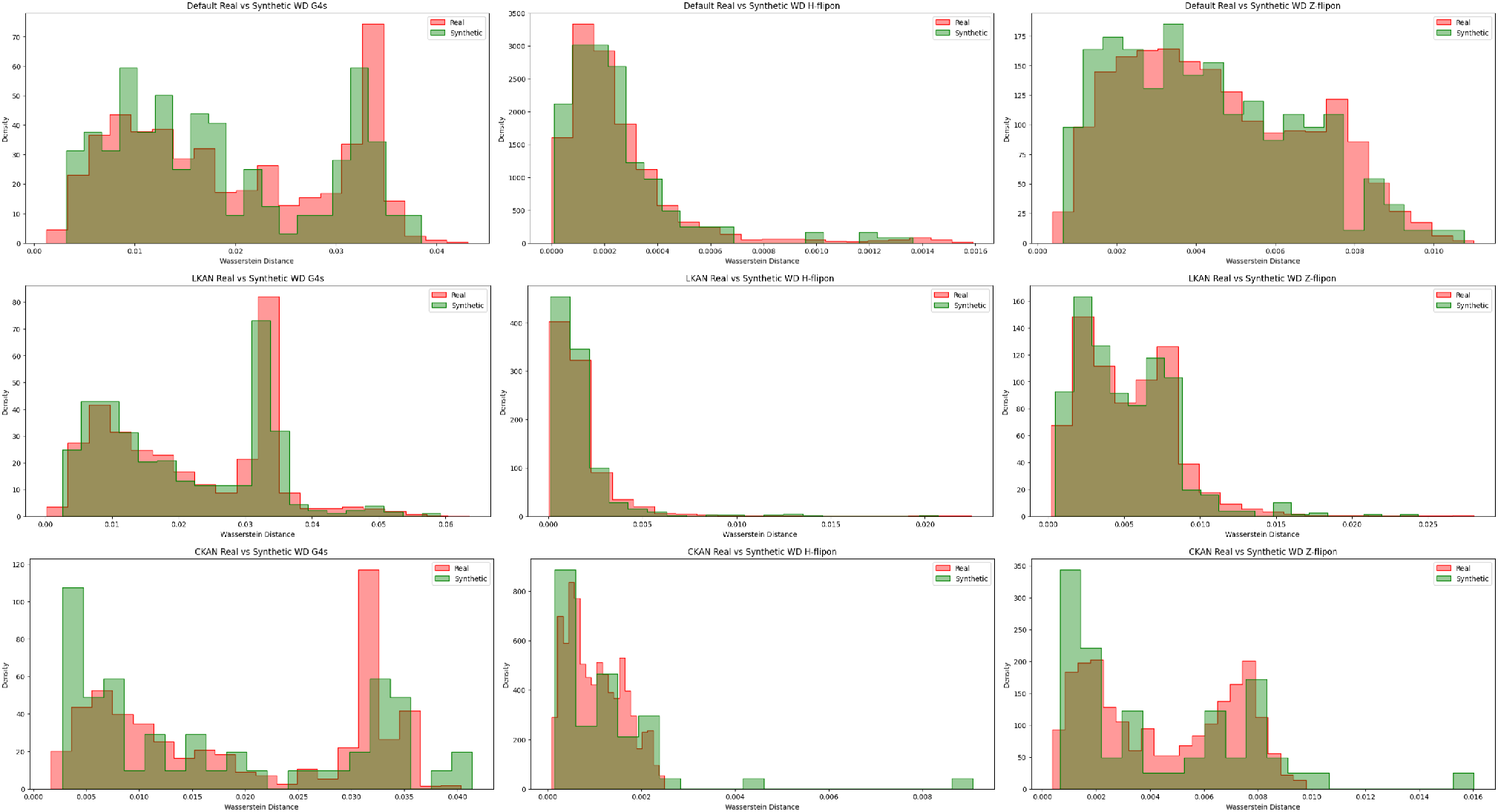
Distributions of Wasserstein Distance (WD) for real and synthetic data. Synthetic data are produced by Baseline Unet (Default) and Unet with Linear KAN (LKAN). We conducted this experiment for flipon benchmark datasets and present distributions for Z-DNA, H-DNA and G4s. Results show that both generative models learned to approximate prior distribution as green and red distributions are close.

In different domains, such as computer vision and natural language processing it is not yet clear if KANs significantly outperform MLPs as it is the topic of active discussions (Yu et al., 2024; Drokin, 2024; Zhou et al., 2024a; Xu et al., 2024b). Our results are preliminary since we have not tested models with large grid size parameters and did not test other types of KANs. Recent advancements in the KAN field inlcude RNN-based and Transformer-based KANs. Temporal Kolmogorov-Arnold Networks (TKAN) using Long Short Term Memory (LSTM) mechanism to leverage time dependency in KANs were proposed in (Genet & Inzirillo, 2024b). Kolmogorov-Arnold Transformer (KAT) architecture is based on Vision Transformer (Dosovitskiy et al., 2021) and utilizes rational polynomial functions to replace splines (Yang & Wang, 2024). KATs were applied to computer vision tasks and, to our knowledge, have not been yet adapted for natural language processing making it difficult for testing on genomic tasks.

Current implementations of Linear KAN lack the same interpretability as original KAN (Liu et al., 2024b) offers: an ability to plot and prune latest layers and suggest symbolic expression for outlying dependencies between variables. Potentially, this could bring even more information about inner structure of a biological task of interest.

Research in the KAN field is evolving rapidly as more and more studies emerge and suggest various technical improvements to address challenges related to KAN usage. The future directions may involve training a Transformer-based models with KANs to handle task of studying latent representations of a language model’s hidden states - embeddings. It was shown (Zhou et al., 2024c; Nguyen et al., 2023) that visualization of language model embeddings could enhance biological understanding of model’s utility. Of interest is to test more frameworks as state-space models (Gu & Dao, 2023), and transformers with GPT-like architecture (Cui et al., 2023). In addition, there are multiple advancements in sequence generative modeling like Discrete Diffusion for DNA (Sarkar et al., 2024). Despite our first positive experiments with linear and convolutional KANs, unleashing the potential of KAN for the entire spectrum of genomic tasks require further extensive research.

## Acknowledgments

The work was supported by the Basic Research Program of the National Research University Higher School of Economics.

