## Supplemental Figure 1 for "Kolmogorov-Arnold Networks for Genomic Tasks"

### 1. SUPPLEMENTARY MATERIALS

#### A. Related Work

In the domain of genomics application of deep learning methods has gained significant traction, particularly with the success of DeepSEA approach [1] - a convolutional neural network designed to predict the chromatin effects of sequence alterations. Subsequently, with the surge of advancements in the field of deep learning, an increasing number of neural networks have been applied to genomics [2–4], [5, 6]. To evaluate deep learning models several studies presented benchmark datasets, which reflect important biological tasks in genomics. One of such studies presented Genomic Benchmarks [7] that is focused on regulatory elements (promoters, enhancers, open chromatin region). Another collection of biological datasets named Genome Understanding Evaluation presented in [8]. The collection includes tasks such as promoter prediction, core promoter prediction, splice site prediction, epigenetic marks prediction, and transcription factor binding sites prediction, and covid variant classification on human and mouse.

Another important genomic task which relates to structural genomics is prediction of non-B (secondary) DNA stuctures such as Z-DNA, G-quadruplexes, H-DNA (triplexes), i-motifs, stem loops, and others. They were termed flipons [9] for they dynamic mode of action. The dynamic nature of non-B formation poses limitations on their discovery at the genome-wide level. The motivation for studying non-B DNA structures stems from their association with various diseases. These structures can influence genomic stability and gene regulation, playing crucial roles in cellular processes. For instance, G-quadruplexes are associated with cancer [10], as they may impact oncogene expression, while Z-DNA is linked to autoimmune diseases due to its involvement in inflammation [11] and immune responses [12]. Existing whole-genome experiments for G4, Z-DNA, H-DNA capture only a subset of all functional flipons in the genome [6, 13–16]. Main challenge one may experience while studying non-B DNA structures is that experimentally obtained data is severely limited.

Current state of the art (SOTA) performance for genomic tasks belongs to Foundation Models such as Large Language Models (LLMs) [17]. For instance, parameter-heavy Transformers such as DNABERT (90M) [18], DNABERT2 (120M) [8], Nucleotide Transformer (500-2500M) [19], GENA-LM (110M) [20], trained on large genomes of several species, are applied to various downstream tasks. In addition, authors of DNABERT2 present Genome Understanding Evaluation (GUE) benchmark - a wide range collection of 28 datasets tackling different genomic tasks. On top of that language models with slightly less amount of parameters that utilize long convolutions - HyenaDNA [21] and Mamba architecture - Caduceus [22] show competitive results with fair performance while significantly reduce time and recourse complexity.

Kolmogorov-Arnold Networks were introduced in [23] as an innovative alternative to traditional MLP by leveraging the Kolmogorov-Arnold representation theorem [24]. Unlike MLP, which employ fixed activation functions on nodes, KANs feature learnable activation functions on edges, replacing linear weight parameters with univariate functions parameterized as splines. However, choice of B-splines raises several challenges in efficient computation, which can be critical aspect when creating deep neural networks with KANs [25]. The following paper [26] presents a comprehensive survey on KANs by incorporating recent studies and advancements. To compare KANs with MLPs in different domains such as computer vision, natural language processing etc, authors of [25] conducted multiple experiments to find out that in original implementation KANs significantly outperforms MLPs only in Symbolic Formula Representing task and in other domains performance is fair. Inspired by approximation in [23] and considering limitations of B-splines, there are multiple works that suggest to use various approximations to replace splines in KAN. One of such works is ReLU-KAN [27], where authors utilize ReLU activation, enabling efficient GPU parallelization. Another study [28] tackles Jacobi basis functions in KANs architecture to speed up training and relatively increase performance compared to original KAN. On top of that, the following work [29] suggest to use a set of radial functions along with B splines to increase training stability. The same goal is pursued by authors of Chebyshev Kolmogorov-Arnold Network [30], where Chebyshev polynomial are placed as basis functions. To address a challenge related to significant time complexity of original KANs when designing deep architectures, authors in [26] present Efficient KAN with pure pytorch implementation.

Inspired by substantial body of research related on Linear KANs [23], two implementations of Convolutional Kolomogorov Arnold Networks (CKANs) have been developped. Primal work [31] presents a novel Convolutional layers based on splines for CKAN. The authors concluded that CKANs maintain top performance compared to original SOTA convolutional neural networks, but use fewer parameters to achieve this performance. They conducted experiments on two standard computer vision datasets such as MNISTs. In [32] authors designed a comprehensive study on convolutional KANs. They utilized a wide range of different convolutional architectures considering various nonlinearity options beyond splines, such as Wavelet transforms and a range of polynomials.

Historically, another type of neural networks - Recurrent Neural Networks (RNNs) has been commonly used for time series abalysis and natural language processing (before the rise of Transformers). This architecture has also been modified by KANs, which resulted in TKAN architecture [33]. Finally, most recent advancement is Kolmogorov Arnold Transformer (KAT) [26], where authors present a Transformer model based on Vision Transformer (ViT) [34] combined with several advancements, such as use of rational polynomial functions to replace splines, to overcome KAN's common limitations.

#### B. Kolmogorov-Arnold Models

We begin by briefly introducing KAN and highlighting the key features of its architecture. Multi-Layer Perceptrons (MLPs) are inspired by the universal approximation theorem [35], which states that a feed-forward network with single hidden layer containing a finite number of neurons can approximate continuous functions on compact subsets of $\mathbb{R}^d$. Kolmogorov-Arnold Network (KAN) focuses on the Kolmogorov-Arnold representation theorem [33], which states that any multivariate continuous function can be represented as a composition of univariate functions and the addition operation

$$f(\mathbf{x}) = \sum_{k=1}^{2n+1} \Phi_k \left( \sum_{l=1}^{n} \phi_{k,l}(x_l) \right) \tag{S1}$$

where $\phi_{k,l}$ are univariates functions that map each input variable $x_l$ such $\phi_{k,l} : [0,1] \to \mathbb{R}$. This means that the (2d+1)(d+1) univariate functions $\Phi_k$ and $\phi_{k,l}$ are enough for an exact representation of a d-variate function. This theorem can be written in matrix form as follows:

$$f(\mathbf{x}) = \Phi_{out} \circ \Phi_{in} \circ \mathbf{x} \tag{S2}$$

Where notations are

$$\begin{aligned} \Phi_{in} &= \begin{bmatrix} \phi_{1,1}(\cdot) & \cdots & \phi_{1,n}(\cdot) \\ \vdots & \ddots & \vdots \\ \phi_{2d+1,1}(\cdot) & \cdots & \phi_{2d+1,d}(\cdot) \end{bmatrix} \\ \Phi_{out} &= \begin{bmatrix} \Phi_1(\cdot), & \ldots, & \Phi_{2d+1}(\cdot) \end{bmatrix} \end{aligned} \tag{S3}$$

This decomposition illustrates how desired function can be built from simpler functions, highlighting a fundamental property of multivariate continuous functions.

A single Kolmogorov-Arnold layer with $d_{in}$-dimensional inputs and $d_{out}$-dimensional outputs in these terms has the following

$$\begin{aligned} f(\mathbf{x}) = \Phi \circ x &= \begin{bmatrix} \sum_{i=1}^{d_{in}} \phi_{i,1}(x_i), & \ldots, & \sum_{i=1}^{d_{in}} \phi_{i,d_{out}}(x_i) \end{bmatrix} \\ \Phi &= \begin{bmatrix} \phi_{1,1}(\cdot) & \cdots & \phi_{1,d_{in}}(\cdot) \\ \vdots & \ddots & \vdots \\ \phi_{d_{out},1}(\cdot) & \cdots & \phi_{d_{out},d_{in}}(\cdot) \end{bmatrix} \end{aligned} \tag{S4}$$

There is also $\Phi = \Phi_{out} \circ \Phi_{in}$. Stacking of $L$ number of layers is represented in a following way

$$\text{KAN}(\mathbf{x}) = (\Phi_{L-1} \circ ... \circ \Phi_1 \circ \Phi_0)\mathbf{x} \tag{S5}$$

Parametrization of $\Phi$ is done by use a linear combination of SiLU activation $\text{SiLU}(\mathbf{x}) = \frac{x}{1+\exp(-x)}$ and a B-spline function $\text{B-spline}(\mathbf{x}) = \sum_i c_i B_i(x)$

$$\phi(\mathbf{x}) = w_b \text{SiLU}(\mathbf{x}) + w_s \text{B-spline}(\mathbf{x}) \tag{S6}$$

The up-to-date paper [36] by the authors of original KAN paper [23] presents several improvements: adapted KAN version for efficient training with gpu, multiplication operation for KANs and an algorithm to compile trained KAN into a network graph to achieve more interpretability.

#### B.1. Kolmogorov Arnold Convolutions

Kolmogorov-Arnold Convolutions could be stated as follows: the kernel consists of a set of univariate non-linear functions. Suppose we have an input image $y \in \mathbb{R}^{c \times h \times w}$, where $c$ is the number of channels, and $h$, $w$ are the height and width of an image respectively. Then, KAN-based convolutions with kernel size $k$ could be defined as

$$x_{ij} = \sum_{d=1}^{c} \sum_{a=0}^{k-1} \sum_{b=0}^{k-1} \varphi_{a,b,d}(y_{d,i+a,j+b}) \tag{S7}$$

Where $i$ in range $[1, h-k+1]$ and $j \in [1, w-k+1]$. In [30] authors describe grid extension as optimization problem

$$\left\{c'_j\right\} = \arg\min_{c'_j} \mathbb{E}_{x \sim p(x)} \left[ \sum_{j=0}^{G_2+k+1} c'_j B(x') - \sum_{j=0}^{G_1+k+1} c_j B(x) \right] \tag{S8}$$

Where $G_1$ is the previous grid size, $G_2$ is the new grid size and $k$ is the B Spline degree. This allows to extend the grid so it gets to be able to capture the variables which escape the original limits.

In [31] authors claim that the main problem with KAN Convolutions lies in the usage of splines, which significantly increase the number of trainable parameters and increase computational cost. They suggest a bottleneck modification of convolutions, which enables a significantly reduction in the number of parameters and scales better. In addition, this study [32] suggests to use Gram polynomial functions along with Radial-Basis Functions, Wavelets, and Chebyshev polynomials. Authors compare these functions to show the advantages of Gram polynomial. Another suggestion presented in [27] is to apply Gaussian noise to a layer's input, thus forcing the model to filter this noise and be more robust against noise in unseen data

$$y'_i = \begin{cases} y_i + \alpha\varepsilon_i, \varepsilon_i \sim \mathcal{N}(0, \sigma^2) \\ y_i \end{cases} \tag{S9}$$

Where $\alpha$ acts as a noise schedule and $\sigma^2$ is a variance of the input, computed for each input channel.

#### B.2. Kolmogorov-Arnold Recurrent Networks.

Authors in [33] proposed Temporal Kolmogorov-Arnold Networks (TKAN) - a novel neural networks architecture inspired by KAN and the LSTM. It utilizes Recurrent Kolmogorov-Arnold Network (RKAN) architecture along with LSTM. We briefly describe Recurring Kolmogorov-Arnold Networks in a way they were presented in [33]. First, authors derive a recurrent kernel in RNNs for state $h_t$ at given time $t$.

$$h_t = f\left(W_{hh} h_{t-1} + W_{hx} x_t + b_h\right) \tag{S10}$$

Where $W_h$ and $W_x$ are the recurrent kernel a weight matrix that transforms the previous hidden states $h_{t-1}$ and the input kernel, a weight matrix transforming the current input denoted $x_t$, respectively. To add positional dependency, authors introduce each transformation $\phi_{l,j,i}$ to be time dependent. Let us denote $h_{l,i}(t)$ a memory function capturing the history of node $i$ in $l$-th layer.

$$x_{l+1,j}(t) = \sum_{i=1}^{n_l} \phi_{l,j,i,t}(x_{l,i}(t), h_{l,i}(t)), j \in [1, n_{l+1}] \tag{S11}$$

Analog for memory step $h_{l,i}(t)$ is defined as a combination of past hidden states

$$h_{l,i}(t) = W_{hh}h_{l,i}(t-1) + W_{hx}x_{l,i}(t) \tag{S12}$$

Rewriting in terms of $\Phi$ matrix

$$x_{l+1}(t) = \begin{bmatrix} \phi_{l,1,1}(\cdot,\cdot) & \dots & \phi_{l,1,n_l}(\cdot,\cdot) \\ \vdots & \ddots & \vdots \\ \phi_{l,n_{l+1},1}(\cdot,\cdot) & \dots & \phi_{l,n_{l+1},n_l}(\cdot,\cdot) \end{bmatrix} \begin{bmatrix} x_l(t) & h_l(t) \end{bmatrix} \tag{S13}$$

Similar to vanilla KAN Layer, Recurring Kolmogorov-Arnold Layer could be expressed as follows

$$\text{RKAN}(x,t) = (\Phi_{L-1,t} \circ \Phi_{L-2,t} \circ \dots \circ \Phi_{0,t})(x,t) \tag{S14}$$

Function that incorporates memory has the following structure

$$\xi(x,t) = \sum_{i_{L-1}=1}^{n_{L-1}} \phi_{L-1,i_L,i_{L-1}} \left( \sum_{i_{L-2}=1}^{n_{L-2}} \dots \left( \sum_{i_0=1}^{n_0} \phi_{0,i_1,i_0}(x_{i_0}(t), h_{0,i_0}(t)) \right) \dots \right) \tag{S15}$$

Next, authors follow LSTM approach and present gates as follows

$$f_t = \sigma(W_f x_t + U_f h_{t-1} + b_f) \tag{S16}$$

Which refers to the forget gate, the input gate

$$i_t = \sigma(W_i x_t + U_i h_{t-1} + b_i) \tag{S17}$$

Output gate

$$o_t = \sigma(\xi(x,t)) \tag{S18}$$

The hidden state, $h_t$, captures the unit's output, while the cell state, $c_t$ is updated such

$$c_t = f_t \odot c_{t-1} + i_t \odot \sigma(W_c x_t + U_c h_{t-1} + b_c) \tag{S19}$$

Predicted output is given by

$$\hat{y}_t = W_{hy}(o_t \odot \tanh(c_t)) + b_y \tag{S20}$$

***B.3. Kolmogorov-Arnold Transformer.***

Authors of Kolmogorov–Arnold Transformer (KAT) [26] replaces the MLPs in vision transformer [34] with KAN layers. They start by given a 2D image $x \in \mathbb{R}^{H \times W \times C}$ flattened to 1D sequence. Then patch embedding and positional encoding are applied to pass it through a series of KAT layers. At layer $l$, the following operations are performed:

$$x_0^{(l)} = \text{MSA}(\text{LN}(x_{l-1})) + x_{l-1}, l = 1, \dots, L \tag{S21}$$

Where MSA refers to Multi-head Self Attention and LN is the layer normalization. For Transformer architecture

$$x_l = \text{MLP}(\text{LN}(x_0^{(l)})) + x_0^{(l)} \tag{S22}$$

Where MLP stands for Multi Layer Perceptron which is replaced by KAT for Kolmogorov Arnold Transformer in a following way

$$x_l = \text{KAT}(\text{LN}(x_0^{(l)})) + x_0^{(l)} \tag{S23}$$

Next, authors of KAT introduce Group-Rational KAN, where they use rational functions as the base function for KAN and share parameters between a group of edges.

$$\phi(x) = wF(x) = \frac{a_0 + a_1 x + \dots + a_m x^m}{1 + |b_1 x + \dots + b_n x^n|} \tag{S24}$$

Authors claim that use of these function enhance the model's expressiveness, stability, and computational efficiency. To describe Group-Rational KAN - share the parameter for the rational function $f$ for each group.

Let $i$ be the index of the input channel. With $g$ groups, each group contains $d_g = d_{in}/g$ channels, where $\lfloor i/d_g \rfloor$ is group index. Then GR-KAN on input vector $x$ can be expressed as

$$\text{GR-KAN}(x) = \Phi \circ x = \left[ \sum_{i=1}^{d_{in}} w_{i,1} F_{\lfloor i/d_g \rfloor}(x_i) \quad \dots \quad \sum_{i=1}^{d_{out}} w_{i,1} F_{\lfloor i/d_g \rfloor}(x_i) \right] \tag{S25}$$

Matrix form looks in a following way

$$\text{GR-KAN}(x) = WF(x) = \begin{bmatrix} w_{1,1} & \dots & w_{1,d_{in}} \\ \vdots & \ddots & \vdots \\ w_{d_{out},1} & \dots & w_{d_{out},d_{in}} \end{bmatrix} \times \left[ F_{\lfloor i/d_g \rfloor}(x_1) \quad \dots \quad F_{\lfloor d_{in}/d_g \rfloor}(x_{d_i n}) \right]^T \tag{S26}$$

In KAT GR-KAN layer as a group-wise rational function F followed by a linear layer

$$\text{GR-KAN}(x) = \text{linear}(\text{GR}(x)) \tag{S27}$$

In this form, sharing parameters across each input channel allows direct application of the rational function to the input vector, equivalently applying it across each grouped edge. In this way, GR-KAN functions as a specialized MLP, with 1) learnable non-linear functions, 2) activation preceding the linear layer, and 3) unique activation functions tailored for each group of edges.

### C. Evaluation metrics of quality

#### *C.1. Metrics for classification*

*Accuracy* measures the proportion of correctly classified instances such as True Positives (TP), True Negatives (TN), False Positives (FP) and False Negatives (FN) out of the total number of instances.

$$\text{ACC} = \frac{\text{TP} + \text{TN}}{\text{TP} + \text{TN} + \text{FP} + \text{FN}} \tag{S28}$$

*Precision*, also known as positive predictive value, gauges the proportion of correctly predicted positive instances among the total predicted positives. It's calculated by considering True Positives against False Positives, following the formula:

$$\text{Precision} = \frac{\text{TP}}{\text{TP} + \text{FP}} \tag{S29}$$

In contrast, *Recall*, also called sensitivity or true positive rate, measures how well the model identifies actual positive instances, crucial when false negatives are costly. Its calculation focuses on TP relative to FN, given by:

$$\text{Recall} = \frac{\text{TP}}{\text{TP} + \text{FN}} \tag{S30}$$

The *F1 Score* is the harmonic mean of precision and recall, giving a balance between the two. It is used when precision and recall are both important, and there is a trade-off between the two.

$$\text{F1} = 2 \times \frac{\text{Recall} \cdot \text{Precision}}{\text{Precision} + \text{Recall}} \tag{S31}$$

*AUC ROC* is an Area Under the Receiver Operating Characteristic curve, which plots the true positive rate (recall) against the false positive rate (FPR) at various threshold settings. The curve shows the trade-off between sensitivity and specificity. *MCC* is Matthews Correlation Coefficient, a balanced measure that takes into account true and false positives and negatives, providing a correlation coefficient between actual and predicted classifications.

$$\text{MCC} = \frac{(\text{TP} \cdot \text{TN}) - (\text{FP} \cdot \text{FN})}{\sqrt{(\text{TP} + \text{FP})(\text{TP} + \text{FN})(\text{TN} + \text{FP})(\text{TN} + \text{FN})}} \tag{S32}$$

##### C.2. Metrics for generation

*Wasserstein distance* is a metric used in Wasserstein Generative Adversarial Networks (WGANs) to measure the distance between the distribution of generated data $\pi(x)$ and the distribution of real data $p(x)$. It provides a more stable and meaningful metric for training GANs than traditional metrics like the Jensen-Shannon divergence used in standard GANs [37].

$$W(p, \pi) = \inf_{\gamma \in \Pi(p,\pi)} \mathbb{E}_{(x,y)\sim\gamma} |x - y| \tag{S33}$$

*Edit distance* or Levenshtein distance is a metric to evaluate similarity (or how close) the sequences are [38].

*KL-Divergence* is Kullback-Leibler Divergence, a measure of how one probability distribution diverges from a second, reference probability distribution.

$$D_{KL}(p, \pi) = \mathbb{E}_{p(x)} \log \left( \frac{p(x)}{\pi(x)} \right) \tag{S34}$$

We use this metric while training diffusion model. To evaluate performance of synthetic sequences we additionally calculate diversity to ensure that sequences within a sample are not the same. Given a dataset $\mathcal{D} = \{(x_0, ..., x_n)\}$ with produced synthetic sequences $x_j$ we calculate Diversity as follows

$$\text{Diversity}(\mathcal{D}) = \frac{1}{|\mathcal{D}|(|\mathcal{D}| - 1)} \sum_{(x_i,y_i)\in\mathcal{D}} \sum_{(x_j,y_j)\in\mathcal{D}\setminus(x_i,y_i)} d(x_i, x_j) \tag{S35}$$

#### D. Interpretation of symbolic representation for genomics.

Despite the various limitations of the original Kolmogorov-Arnold Networks [23] such as the computational and time complexity of splines, a significant contribution of original KAN lies in the ability to achieve interpretability through symbolic representation. The authors suggested the following algorithm.

- Start by training a fully-connected KAN model with sparsification regularization; this can result in a significantly sparse architecture.
- Automatic pruning discard all hidden neurons except the last one, leaving one neuron KAN. The activation functions appears to be known symbolic functions. Use suggested symbolic expression as output of KAN model.
- After symbolifying all the activation functions in the network, continue training these affine parameters, and when we see the loss dropping to machine precision, we know that we have found the correct symbolic expression.
- Output the symbolic formula and plot the model.

To investigate a potential of this method we train one layer of KAN using Drosophila enhancers Stark dataset from Genomic Benchmark collection.

To our best knowledge, at the current moment implementations of advanced KAN models such as ConvKANs and Kolmogorov Arnold Transformer (KAT) do not utilize pruning and symbolic representation, which makes such way of interpretation challenging for a wide range of complicated datasets where Vanilla KANs show poor performance.

#### E. Models

Here we present more details on various architectures we used in our work. Overall, we used seven different models. For classification task we utilize the following models: LegNet based convolutional neural network as our baseline model (Baseline), Linear Kolmogorov Arnold Network (LKAN) where we changed MLP layers in baseline model with linear KAN, Convolutional Kolmogorov Arnold Network (CKAN) where we changed convolutional layers in EfficientNet [39] block of LegNet. In Ablation study we present experiment on how changing layers impact the performance. For generative task we used the following models: Denoising Unet based on DNA-Diffusion framework (Baseline), similar to classification changes with Linear KAN where we changed MLPs with LKANs and exactly the same for CKAN. We do not provide the results for combination of LKAN and CKAN for our experiments due to resource limitaitions (see Limitations). Below we present detailed schemes on how we changed baseline models. We trained our models for classification with AdamW optimizer ($\alpha = 5 \cdot 10^{-4}, w = 5 \cdot 10^{-3}$) with tested OneCycleLR. Since datasets we used already split in train/test we also tested cross validation with 5 folds for training data and then applied trained model for separate evaluation. For generative models we used Exponential Moving Average (EMA) and gradient accumulation with size of 8. The normalization on convolution blocks is made by GroupNorm layers and SiLU instead of GeLU. The most stable training was observed under AdamW optimizer ($\alpha = 10^{-4}, \beta_1 = 0.99, \beta_2 = 0.99, w = 5 \cdot 10^{-3}$). For the loss we considered three options: l1, l2 and huber loss function. Last option allowed us to get faster convergence of losses.

#### F. Flipon datasets

**Motivation** Flipons are important functional genetic elements that regulate various genomic processes by changing conformation. Recent studies highlighted the key-role of flipons in immune response, either switching interferon responses of to limit infammation or initiating cell death to eliminate virally infected or dysfunctional cells. Te fipons involved are subject to natural selection and underscore the importance of fipons in genome evolution.

**Collection** To create datasets for nonB DNA structures we considered the following experiments [40–43]. Selected datasets are divided into three categories: for G-quadruplexes (G4s): Endoquad, G4-ChIP, G4-seq and G4 CUT&Tag; for Z-DNA: [44, 45]; for H-DNA: [44]. Sequences were obtained by alignment to reference genome hg19. Negative samples were selected from the same genome as the positive samples. For each positive sample, we generated a random interval in the genome with the same length as a given sample. We picked only those intervals not overlapping with any of the positive samples.

**Composition** Full description of datasets is presented at Supplementary Table 1.

*Endoquad* has 280,532 sequences with a median length of 32 base pairs and a standard deviation of 8.6. The dataset is split into two equal classes (class ratio 1.0) with a 7:3 train/test ratio.

*G4-ChIP* dataset contains 8,900 sequences with a median length of 128 base pairs and a larger standard deviation of 35.9, also maintaining a 7:3 train/test ratio and equal class distribution.

*G4 CUT&Tag* experiment has 17,880 sequences, a median length of 206 base pairs, and a higher variability with a standard deviation of 118.6, with the same 7:3 train/test split.

*G4-seq* is the largest dataset with 1,402,000 sequences, a median length of 137 base pairs, and a standard deviation of 56.9, with an equal class ratio and a 7:3 split.

*Kouzine Z-DNA* comprises 85,308 sequences with a median length of 23 base pairs and a standard deviation of 26.2, also using a 7:3 split.

*Shin Z-DNA* has only 726 sequences, with a much longer median sequence length of 384 base pairs and a standard deviation of 98.4, with an 8:2 train/test split.

*Kouzine H-DNA* dataset includes 1,409,662 sequences with a median length of 28 base pairs and a standard deviation of 25.4, maintaining a 7:3 train/test split.

Across all datasets, the class ratio remains consistent at 1.0, indicating balanced classes in all cases. The datasets vary significantly in the number of sequences, sequence lengths, and variability, offering a diverse set of non-B DNA data for classification tasks.

#### G. Limitations

Our results are preliminary since we have not tested models with large grid size parameter limited to our computational resources. Moreover, we did not test some of the recent architectures in KAN's field like Kolmogorov Arnold Transformer (KAT) [26] since they are not yet adapted for natural language processing. Research toward KANs is rapidly changing as more and more research papers emerge and suggest various technical improvements to address challenges with KANs. In this work we aimed to test more popular KAN implementations like Linear KAN and Convolutional KAN for handling genomic data. To accomplish this task we tried to use a wide variety of datasets with different biological tasks. However, in terms of machine learning impact our research is shifted to sequence generation and classification. To bring more insights, our future plans involve training Transformer-based models with KANs to handle task of studying latent representations of language model's hidden states - embeddings. It has been shown [21, 46] that visualization of language model's embeddings could enhance biological understanding of model's impact.

Another limitation of our research is that we could not test larger sizes for our models and could not conduct corresponding experiments in parallel as we utilize only single A100 gpu via colab. This tremendously shifts us in terms scaling of our models. For instance, when trying to combine Linear KAN and Convolutional KAN together we faced a challenge to maintain the same performance and scale grid size parameter which leads to longer training process.
Lastly, current implementations of Linear KAN lacks the same interpretability that original KAN [23] offers - an ability to plot and prune latest layers and suggest symbolic expression for outlying dependencies between variables. This could bring even more information about inner structure of biological task of interest.

As our future plans, we aim to extend impact on meaningful genomic tasks collecting more datasets and specific machine learning tasks. Moreover, we plan to test more frameworks as state space models [47], and transformers with GPT like architecture [48]. In addition, there are multiple advancements in sequence generative modeling like Discrete Diffusion for DNA [49].

#### H. Extended experimental results

Here we present extended results of conducted experiments for classification and generation of DNA sequences for each of the collection we follow in our research: Genomic Benchmarks (GB), Genome Understanding Evaluation (GUE), extended version of Genome Understanding Evaluation(GUE+). We construct extended results as follows, say compare baseline model (model presented in original paper of each dataset's collection) and state of the art (model that achieves the highest score for the collection) with a variety of KAN based models - Vanilla KAN (VKAN), Linear KAN (LKAN) and Convolutional KAN (CKAN). For GUE we also compare KAN's performance with HyenaDNA along with CNN model from [7] and [8].

For extended results in Genomic Benchmarks given the equal size of models we observe that LKAN achieves the highest performance, overall outperforming baseline CNN, yet rarely achieving SOTA result of HyenaDNA. For flipon's collection of datasets we observe that both LKAN and CKAN outperform baseline models performing close with SOTA language models.

Results are given in Supplementary Tables 3-4. For Genome Understanding Evaluation (both GUE and GUE+) LKAN significantly outperforms baseline CNN and often outperforms HyenaDNA, yet keeps getting far from DNABERT2. The results are presented in Tables 2-14.

For generative models we present extended results for evaluation of grid size impact. As we showed earlier increasing grid size in Linear KAN significantly increases amount of model's parameters. Hence, it makes the training process more challenging in terms of resources. However, the upside of this modification is a positive trend for increasing the performance. We utilize Linear KAN layers and 2d Convolutional KAN layers in Unet architecture and observe that LKANs scales better with grid size than CKANs that is shown at Figure 1a. For instance, when evaluating generative models with KL divergence and Wasserstein distance, we observe that to achieve the same performance LKAN needs grid size 10 (total number of parameters equals 200M) while CKAN requires grid size of 30 (total number of parameters 800M).

### 2. SUPPLEMENTARY FIGURES AND TABLES

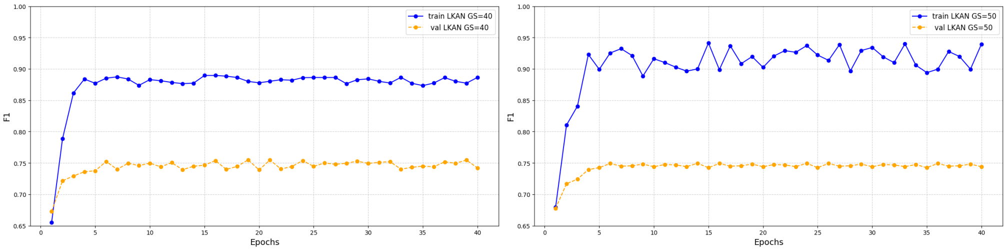


**Fig. S1.** Comparison of LKAN models with high values of grid size, 40 and 50 respectively, to demonstrate overfit. Notably, runs with higher grid size results in spiky training curves.

**Table S1.** Description of flipons datasets

| Dataset | Num of seqs | Num | Class ratio | Median length | Std | Train/Test |
|---|---|---|---|---|---|---|
| EndoQuad | 280532 | 2 | 1.0 | 32.0 | 8.6 | 7:3 |
| G4ChIP | 8900 | 2 | 1.0 | 128.0 | 35.9 | 7:3 |
| G4cut | 17880 | 2 | 1.0 | 206.0 | 118.6 | 7:3 |
| G4seq | 1402000 | 2 | 1.0 | 137 | 56.9 | 7:3 |
| Kou | 85308 | 2 | 1.0 | 23.0 | 26.2 | 7:3 |
| Shin | 726 | 2 | 1.0 | 384.0 | 98.4 | 8:2 |
| HKou | 1409662 | 2 | 1.0 | 28.0 | 25.4 | 7:3 |

**Table S2.** Classification results for Genomic Benchmarks.

| Model | Acc | AUC | PR | F1 | MCC | Better? |
|---|---|---|---|---|---|---|
| **Dummy Mouse Enhancers Ensembl - DMEE** | | | | | | |
| Default CNN | 69.0± 0.5 | 75.2± 0.4 | 68.3± 0.1 | 70.4± 0.1 | 49.4± 0.4 | √ |
| HyenaDNA (SOTA) | 85.1± 0.5 | 86.2± 0.4 | 80.2± 0.7 | 82.3± 0.4 | 79.9± 0.5 | × |
| Baseline LegNet | 80.5± 0.6 | 84.4± 0.5 | 78.7± 0.8 | 82.1 ± 0.5 | 70.2± 0.5 | √ |
| LKAN | 84.0± 0.7 | 85.5± 0.6 | 79.6± 0.9 | 84.0± 0.4 | 73.2± 0.2 | √ |
| CKAN | 81.1± 0.9 | 83.3± 0.9 | 77.7± 1.2 | 78.1± 0.6 | 64.8± 1.3 | √ |
| **Demo Coding vs Intergenomic Seqs - DCIS** | | | | | | |
| Default CNN | 87.6± 0.7 | 88.3± 0.3 | 84.3± 0.2 | 86.8± 0.7 | 80.2± 0.6 | √ |
| HyenaDNA (SOTA) | 91.3± 0.5 | 93.0± 0.9 | 88.3± 0.1 | 90.4± 0.9 | 87.9± 0.6 | × |
| Baseline LegNet | 89.9± 0.5 | 92.3± 0.5 | 87.0± 0.5 | 88.4± 0.8 | 81.1± 0.5 | √ |
| LKAN | 90.6± 0.4 | 93.1± 0.4 | 87.3± 0.5 | 91.2± 0.7 | 85.5± 1.4 | √ |
| CKAN | 85.9± 1.2 | 85.0± 1.2 | 85.3± 1.2 | 88.7± 1.2 | 81.7± 1.2 | √ |
| **Demo Human or Worm - DHW** | | | | | | |
| Default CNN | 93.0± 0.2 | 91.8± 0.1 | 88.2± 0.2 | 92.8± 0.5 | 89.2± 0.5 | √ |
| HyenaDNA (SOTA) | 96.6± 0.4 | 96.5± 0.1 | 92.3± 0.4 | 94.7± 0.6 | 90.2± 0.4 | × |
| Baseline LegNet | 94.4± 0.5 | 92.2± 0.7 | 90.0± 0.5 | 93.9± 0.5 | 89.2± 0.5 | √ |
| LKAN | 95.1± 0.6 | 95.9± 0.5 | 91.1± 0.4 | 94.4± 0.5 | 90.2± 0.5 | √ |
| CKAN | 94.9± 1.0 | 94.5± 0.7 | 90.4± 0.5 | 92.1± 0.8 | 88.1± 0.8 | √ |
| **Drosophila Enhancers Stark - DES** | | | | | | |
| Default CNN | 58.6± 0.1 | 56.8± 0.4 | 55.5± 0.1 | 44.5± 0.4 | 43.2± 0.6 | √ |
| HyenaDNA (SOTA) | 58.6± 0.4 | 58.8± 0.3 | 53.7± 0.5 | 50.3± 0.1 | 49.4± 0.7 | × |
| Baseline LegNet | 58.0± 0.4 | 56.5± 0.6 | 45.2± 0.2 | 44.8± 0.7 | 27.3± 0.9 | √ |
| LKAN | 58.4± 0.1 | 59.9± 0.3 | 49.4± 0.7 | 44.5± 0.5 | 28.9± 0.6 | √ |
| CKAN | 57.5± 0.3 | 56.6± 0.4 | 47.1± 0.7 | 43.2± 0.8 | 25.3± 0.6 | √ |
| **Human Enhancers cohn - HEC** | | | | | | |
| Default CNN | 69.5± 0.4 | 70.0± 0.1 | 67.3± 0.4 | 67.1± 0.5 | 63.6± 0.1 | √ |
| HyenaDNA (SOTA) | 74.2± 0.1 | 72.3± 0.3 | 70.8± 0.4 | 71.2± 0.4 | 70.0± 0.4 | × |
| Baseline LegNet | 73.3± 0.5 | 72.0± 0.4 | 68.8± 0.5 | 70.1± 0.7 | 66.6± 0.8 | √ |
| LKAN | 74.0± 0.5 | 72.0± 0.2 | 69.0± 0.4 | 73.4±0.7 | 68.2± 0.5 | √ |
| CKAN | 73.0± 0.9 | 71.4± 0.8 | 67.6± 0.7 | 70.7± 0.8 | 63.9± 0.8 | √ |
| **Human Enhancers Ensembl - HEE** | | | | | | |
| Default CNN | 68.9± 0.5 | 70.1± 0.4 | 57.3± 0.5 | 56.5± 0.4 | 55.3± 0.4 | √ |
| HyenaDNA (SOTA) | 89.2± 0.3 | 91.1± 0.5 | 87.4± 0.6 | 89.8± 0.5 | 86.5± 0.3 | × |
| Baseline LegNet | 75.5± 0.7 | 76.6± 0.6 | 64.6± 0.6 | 60.5± 0.7 | 63.2± 0.5 | √ |
| LKAN | 80.2± 0.8 | 83.3± 1.1 | 74.4± 0.9 | 67.2± 0.8 | 69.9± 0.5 | √ |
| CKAN | 75.5± 1.0 | 80.2± 1.0 | 67.7± 1.0 | 58.3± 0.8 | 59.0± 1.0 | √ |

**Table S3.** Classification results for Genomic Benchmarks.*(Continued from previous page)*.

| Human Ensembl Regulatory - HER | | | | | | |
|---|---|---|---|---|---|---|
| Model | Acc | AUC | PR | F1 | MCC | Better? |
| Default CNN | 93.3± 0.1 | 94.2± 0.4 | 90.2± 0.6 | 93.3± 0.3 | 89.2± 0.8 | √ |
| HyenaDNA (SOTA) | 93.8± 0.4 | 95.5± 0.5 | 94.4± 0.6 | 93.6± 0.5 | 91.1± 0.1 | × |
| Baseline LegNet | 93.5± 0.7 | 94.0± 0.7 | 91.1± 0.7 | 93.5± 0.7 | 90.0± 0.8 | √ |
| LKAN | 93.5± 0.8 | 91.9± 0.6 | 90.7± 0.8 | 93.2± 0.7 | 90.3± 0.8 | √ |
| CKAN | 93.0± 0.6 | 93.5± 0.5 | 89.4± 0.7 | 92.4± 0.5 | 89.4± 0.6 | √ |
| Human Nontata Promoters - HNP | | | | | | |
| Default CNN | 84.6± 0.4 | 86.6± 0.5 | 82.4± 0.4 | 83.7± 0.5 | 81.1± 0.1 | √ |
| HyenaDNA (SOTA) | 96.6± 0.3 | 94.4± 0.7 | 94.3± 0.5 | 95.0± 0.5 | 90.2± 0.4 | × |
| Baseline LegNet | 91.6± 0.7 | 92.3± 0.7 | 90.0± 0.5 | 90.2± 0.8 | 87.9± 0.9 | √ |
| LKAN | 95.0± 0.7 | 93.9± 0.6 | 91.6± 0.7 | 94.5± 0.9 | 88.8± 0.9 | √ |
| CKAN | 92.0± 1.2 | 89.7± 1.1 | 87.0± 1.0 | 88.8± 1.1 | 85.0± 1.0 | √ |
| Human Ocr Ensembl - HOE | | | | | | |
| Default CNN | 68.0± 0.1 | 67.7± 0.3 | 63.1± 0.1 | 66.1± 0.3 | 60.1± 0.3 | √ |
| HyenaDNA (SOTA) | 80.9± 0.5 | 83.3± 0.7 | 81.1± 0.4 | 82.4± 0.4 | 78.5± 0.5 | √ |
| Baseline LegNet | 76.6± 0.6 | 80.3 ± 0.7 | 75.0± 0.9 | 74.5± 0.5 | 70.8± 0.6 | √ |
| LKAN | 78.4± 0.5 | 79.6± 0.5 | 78.0± 0.6 | 79.2± 0.8 | 74.4± 0.7 | √ |
| CKAN | 73.6± 1.0 | 75.5± 1.0 | 69.0± 1.0 | 72.7± 1.0 | 68.9± 1.0 | √ |

**Table S4.** Classification results for flipons.

| ENDOQUAD | | | | | | |
|---|---|---|---|---|---|---|
| MODEL | ACC | AUC | PR | F1 | MCC | BETTER? |
| DEEPZ | 86.8± 0.7 | 89.4± 0.5 | 84.5± 0.6 | 88.5± 0.5 | 80.3± 0.6 | √ |
| DNABERT2 (SOTA) | 97.4± 0.5 | 96.0± 0.5 | 98.4± 0.5 | 97.7± 0.4 | 97.2± 0.5 | × |
| BASELINE LEGNET | 95.1± 0.6 | 94.8± 0.6 | 93.7± 0.6 | 92.9± 0.2 | 89.1± 0.5 | √ |
| LKAN | 95.9± 0.4 | 96.0± 0.5 | 92.6± 0.5 | 96.0± 0.2 | 93.5± 0.4 | √ |
| CKAN | 92.4± 0.8 | 93.0± 0.8 | 88.5± 0.8 | 89.8± 0.8 | 89.0± 0.8 | √ |
| **G4SEQ** | | | | | | |
| DEEPZ | 83.1± 0.5 | 84.7± 0.5 | 80.1± 0.7 | 83.0± 0.7 | 77.7± 0.7 | √ |
| DNABERT2 (SOTA) | 95.6± 0.3 | 96.1± 0.3 | 94.0± 0.3 | 97.1± 0.3 | 95.0± 0.3 | × |
| BASELINE LEGNET | 86.0± 0.6 | 87.3± 0.6 | 84.2± 0.6 | 87.5± 0.6 | 83.3± 0.6 | √ |
| LKAN | 92.3± 0.7 | 91.6± 0.7 | 89.6± 0.7 | 89.4± 0.5 | 89.0± 0.8 | √ |
| CKAN | 85.9± 0.2 | 84.6± 0.5 | 85.7± 0.4 | 85.4± 0.4 | 84.5± 0.2 | √ |
| **G4CHIP** | | | | | | |
| DEEPZ | 93.5± 0.5 | 94.1± 0.5 | 88.1± 0.5 | 90.9± 0.5 | 88.9± 0.5 | √ |
| DNABERT2 (SOTA) | 99.1± 0.4 | 98.5± 0.4 | 98.8± 0.4 | 97.9± 0.4 | 97.0± 0.6 | × |
| BASELINE LEGNET | 95.0± 0.6 | 96.4± 0.5 | 93.2± 0.5 | 91.1± 0.5 | 90.2± 0.9 | √ |
| LKAN | 97.3± 0.4 | 96.8± 0.7 | 93.4± 0.5 | 93.0± 0.4 | 92.2± 0.7 | √ |
| CKAN | 94.6± 1.0 | 95.0± 1.0 | 90.2± 1.1 | 94.8± 1.2 | 91.3± 1.5 | √ |
| **G4CUT** | | | | | | |
| DEEPZ | 91.2± 0.5 | 93.0± 0.5 | 88.7± 0.5 | 90.3± 0.6 | 89.5± 0.5 | √ |
| DNABERT2 (SOTA) | 95.4± 0.5 | 97.8± 0.5 | 96.4± 0.5 | 96.1± 0.5 | 96.0± 0.5 | × |
| BASELINE LEGNET | 93.0± 0.5 | 94.7± 0.6 | 90.8± 0.5 | 92.6± 0.7 | 91.1± 0.8 | √ |
| LKAN | 95.0± 0.7 | 95.5± 0.7 | 93.3± 0.7 | 96.0± 0.6 | 94.8± 0.7 | √ |
| CKAN | 91.5± 1.0 | 92.4± 1.0 | 87.0± 1.0 | 91.4± 1.0 | 85.6± 1.2 | √ |

**Table S5.** Classification results for flipons. *(Continued from previous page)*

| KOU | | | | | | |
|---|---|---|---|---|---|---|
| Model | Acc | AUC | PR | F1 | MCC | Better? |
| DeepZ | 89.0± 0.7 | 88.4± 0.6 | 85.0± 0.5 | 87.7± 0.5 | 85.3± 0.5 | √ |
| DNABERT2 (SOTA) | 98.0± 0.4 | 99.1± 0.4 | 97.4± 0.4 | 98.5± 0.4 | 97.0± 0.4 | √ |
| Baseline LegNet | 93.0± 1.0 | 95.2± 1.0 | 91.3± 1.0 | 95.7± 1.0 | 94.5± 1.0 | √ |
| LKAN | 96.5± 0.7 | 94.8± 0.7 | 94.8± 0.6 | 97.1± 0.8 | 96.3± 0.8 | √ |
| CKAN | 91.3± 0.5 | 94.3± 0.7 | 91.0± 0.7 | 95.6± 0.7 | 94.0± 0.8 | √ |
| **Shin** | | | | | | |
| DeepZ | 95.4± 0.3 | 95.0± 0.3 | 95.9± 0.3 | 95.9± 0.3 | 94.5± 0.3 | √ |
| DNABERT2 (SOTA) | 99.0± 0.1 | 99.5± 0.1 | 99.2± 0.1 | 99.1± 0.1 | 98.9± 0.1 | √ |
| Baseline LegNet | 98.5± 0.5 | 97.0± 0.5 | 96.7± 0.5 | 98.9± 0.5 | 97.9± 0.5 | × |
| LKAN | 98.0± 0.6 | 96.6± 0.5 | 97.0± 0.5 | 96.5± 0.5 | 97.0± 0.5 | √ |
| CKAN | 95.3± 1.0 | 96.0± 1.0 | 93.0± 1.0 | 97.4± 1.0 | 95.7± 1.1 | √ |
| **HKOU** | | | | | | |
| DeepZ | 80.4± 0.5 | 82.1± 0.5 | 79.5± 0.6 | 82.3± 0.5 | 78.9± 0.8 | √ |
| DNABERT2 (SOTA) | 97.0± 0.4 | 96.1± 0.4 | 96.5± 0.4 | 98.3± 0.4 | 96.6± 0.4 | √ |
| Baseline LegNet | 93.1± 0.6 | 92.6± 0.62 | 89.4± 0.6 | 93.4± 0.6 | 90.5± 0.6 | √ |
| LKAN | 95.0± 0.9 | 93.1± 0.9 | 90.4± 0.9 | 96.4± 0.9 | 95.2± 1.1 | √ |
| CKAN | 94.3± 0.9 | 90.0± 0.9 | 85.5± 0.9 | 90.8± 0.9 | 88.7± 0.9 | √ |

**Table S6.** Classification results for GUE

| Epigenetic Marks Prediction | | | | | | |
|---|---|---|---|---|---|---|
| | | | *H3* | | | |
| Model | Acc | AUC | PR | F1 | MCC | Better? |
| Baseline CNN | 65.4± 0.7 | 66.6± 0.7 | 63.5± 0.7 | 60.0± 0.7 | 61.5± 0.7 | √ |
| DNABERT2 (SOTA) | 79.4± 0.4 | 82.0± 0.4 | 77.4± 0.4 | 81.5± 0.4 | 78.3± 0.4 | × |
| HyenaDNA | 70.0± 0.5 | 74.5± 0.5 | 67.0± 0.5 | 71.0± 0.5 | 67.2± 0.5 | √ |
| Baseline LegNet | 68.0± 0.8 | 70.5± 0.7 | 65.0± 0.7 | 71.5± 0.7 | 70.7± 1.5 | √ |
| LKAN | 71.5± 0.6 | 75.0± 0.5 | 68.0± 0.6 | 72.0± 0.6 | 71.3± 1.0 | √ |
| CKAN | 67.8± 0.7 | 70.0± 0.6 | 64.5± 0.6 | 66.2± 0.6 | 69.4± 1.5 | √ |
| | | | *H3K14ac* | | | |
| Baseline CNN | 50.4± 0.5 | 55.3± 0.2 | 29.0± 0.2 | 33.2± 0.2 | 29.7± 0.2 | √ |
| DNABERT2 (SOTA) | 72.3± 0.4 | 70.9± 0.4 | 55.5± 0.4 | 60.7± 0.4 | 52.6± 0.4 | × |
| HyenaDNA | 60.0± 0.6 | 61.3± 0.6 | 35.0± 0.6 | 40.7± 0.6 | 32.0± 0.6 | √ |
| Baseline LegNet | 54.0± 0.6 | 57.0± 0.5 | 31.5± 0.5 | 35.0± 0.5 | 40.2± 0.8 | √ |
| LKAN | 58.5± 0.5 | 62.0± 0.5 | 37.0± 0.5 | 42.0± 0.5 | 43.2± 0.7 | √ |
| CKAN | 53.5± 0.6 | 56.5± 0.6 | 30.5± 0.6 | 34.5± 0.6 | 41.0± 1.0 | √ |
| | | | *H3K36me3* | | | |
| Baseline CNN | 56.6± 0.5 | 58.1± 0.5 | 45.6± 0.5 | 58.0± 0.5 | 38.6± 0.5 | √ |
| DNABERT2 (SOTA) | 75.6± 0.5 | 75.7± 0.5 | 68.4± 0.5 | 70.8± 0.5 | 56.9± 0.5 | × |
| HyenaDNA | 67.0± 0.4 | 63.2± 0.4 | 60.1± 0.4 | 70.2± 0.4 | 48.3± 0.4 | √ |
| Baseline LegNet | 59.5± 0.6 | 60.5± 0.6 | 48.5± 0.6 | 59.2± 0.6 | 44.3± 0.9 | √ |
| LKAN | 63.5± 0.5 | 65.0± 0.5 | 52.5± 0.5 | 64.0± 0.5 | 47.7± 0.6 | √ |
| CKAN | 58.7± 0.6 | 59.8± 0.6 | 47.8± 0.6 | 58.0± 0.6 | 42.9± 0.9 | √ |
| | | | *H3K4me1* | | | |
| Baseline CNN | 55.1± 0.1 | 56.3± 0.1 | 37.0± 0.1 | 35.8± 0.1 | 26.1± 0.1 | √ |
| DNABERT2 (SOTA) | 75.0± 0.3 | 76.4± 0.3 | 69.0± 0.3 | 71.3± 0.3 | 50.5± 0.3 | × |
| HyenaDNA | 65.9± 0.5 | 67.3± 0.5 | 50.1± 0.5 | 58.4± 0.5 | 35.9± 0.5 | √ |
| Baseline LegNet | 57.2± 0.4 | 58.0± 0.4 | 38.5± 0.4 | 37.0± 0.4 | 35.6± 0.4 | √ |
| LKAN | 66.5± 0.5 | 67.8± 0.5 | 52.5± 0.5 | 61.5± 0.5 | 40.0± 0.8 | √ |
| CKAN | 56.5± 0.6 | 57.5± 0.6 | 38.2± 0.6 | 36.5± 0.6 | 35.8± 1.1 | √ |
| | | | *H3K4me2* | | | |
| Baseline CNN | 57.8± 0.4 | 50.5± 0.4 | 28.9± 0.4 | 40.6± 0.4 | 25.8± 0.4 | √ |
| DNABERT2 (SOTA) | 67.2± 0.3 | 66.6± 0.3 | 39.0± 0.3 | 51.0± 0.3 | 31.1± 0.3 | × |
| HyenaDNA | 58.0± 0.5 | 55.6± 0.5 | 35.6± 0.5 | 47.7± 0.5 | 25.8± 0.5 | √ |
| Baseline LegNet | 59.0± 0.6 | 52.3± 0.6 | 30.1± 0.6 | 42.2± 0.6 | 27.2± 0.5 | √ |
| LKAN | 60.5± 0.5 | 54.0± 0.5 | 32.5± 0.5 | 44.8± 0.5 | 29.3± 0.5 | √ |
| CKAN | 57.5± 0.7 | 50.8± 0.7 | 29.2± 0.7 | 40.1± 0.7 | 24.3± 1.2 | √ |
| | | | *H3K4me3* | | | |
| Baseline CNN | 54.6± 0.3 | 54.8± 0.3 | 30.9± 0.3 | 41.4± 0.3 | 20.5± 0.3 | √ |
| DNABERT2 (SOTA) | 63.0± 0.4 | 65.1± 0.4 | 44.4± 0.4 | 55.0± 0.4 | 36.3± 0.4 | × |
| HyenaDNA | 57.0± 0.5 | 57.8± 0.5 | 35.4± 0.5 | 43.3± 0.5 | 23.2± 0.5 | √ |
| Baseline LegNet | 55.5± 0.6 | 56.0± 0.6 | 31.5± 0.6 | 42.0± 0.6 | 28.4± 0.5 | √ |
| LKAN | 58.0± 0.5 | 58.6± 0.5 | 33.9± 0.5 | 45.0± 0.5 | 29.8± 0.4 | √ |
| CKAN | 54.8± 0.7 | 55.2± 0.7 | 30.7± 0.7 | 41.2± 0.7 | 27.2± 0.7 | √ |

**Table S7.** Classification results for GUE *(Continued from previous page)*.

| Epigenetic Marks Prediction | | | | | | |
|---|---|---|---|---|---|---|
| *H3K79me3* | | | | | | |
| Model | Acc | AUC | PR | F1 | MCC | Better? |
| Baseline CNN | 65.9± 0.2 | 65.9± 0.2 | 45.4± 0.2 | 60.2± 0.2 | 46.3± 0.2 | √ |
| DNABERT2 (SOTA) | 84.6± 0.5 | 85.0± 0.5 | 80.0± 0.5 | 78.4± 0.5 | 67.4± 0.5 | × |
| HyenaDNA | 71.7± 0.3 | 73.2± 0.3 | 55.4± 0.3 | 64.5± 0.3 | 54.1± 0.3 | √ |
| Baseline LegNet | 68.5± 0.4 | 67.8± 0.4 | 50.0± 0.4 | 61.0± 0.4 | 60.8± 0.9 | √ |
| LKAN | 70.2± 0.5 | 69.4± 0.5 | 52.5± 0.5 | 63.5± 0.5 | 62.1± 0.8 | √ |
| CKAN | 68.9± 0.6 | 67.9± 0.6 | 50.8± 0.6 | 61.5± 0.6 | 61.1± 1.5 | √ |
| *H3K9ac* | | | | | | |
| Baseline CNN | 61.0± 0.5 | 59.8± 0.5 | 55.4± 0.5 | 52.7± 0.5 | 40.0± 0.5 | √ |
| DNABERT2 (SOTA) | 70.1± 0.2 | 73.9± 0.2 | 65.5± 0.2 | 60.4± 0.2 | 55.6± 0.5 | × |
| HyenaDNA | 65.4± 0.4 | 66.6± 0.4 | 58.9± 0.4 | 61.0± 0.4 | 50.8± 0.4 | √ |
| Baseline LegNet | 63.5± 0.5 | 61.9± 0.5 | 56.8± 0.5 | 54.2± 0.5 | 50.5± 0.6 | √ |
| LKAN | 65.0± 0.6 | 63.5± 0.6 | 59.0± 0.6 | 56.5± 0.6 | 51.9± 0.5 | √ |
| CKAN | 64.0± 0.7 | 62.5± 0.7 | 57.5± 0.7 | 55.0± 0.7 | 50.0± 0.9 | √ |
| *H4* | | | | | | |
| Model | Acc | AUC | PR | F1 | MCC | Better? |
| Baseline CNN | 81.3± 0.7 | 80.7± 0.7 | 78.7± 0.7 | 70.8± 0.7 | 62.3± 0.7 | √ |
| DNABERT2 (SOTA) | 90.5± 0.5 | 92.6± 0.5 | 86.6± 0.5 | 85.0± 0.5 | 80.7± 0.5 | × |
| HyenaDNA | 86.8± 0.4 | 85.9± 0.4 | 81.1± 0.4 | 79.5± 0.4 | 73.7± 0.4 | × |
| Baseline LegNet | 83.0± 0.6 | 82.2± 0.6 | 79.0± 0.6 | 73.4± 0.6 | 75.6± 0.9 | √ |
| LKAN | 84.2± 0.6 | 83.5± 0.6 | 80.2± 0.6 | 75.2± 0.6 | 77.2± 0.7 | √ |
| CKAN | 82.5± 0.7 | 81.8± 0.7 | 78.5± 0.7 | 72.8± 0.7 | 70.9± 0.8 | √ |
| *H4ac* | | | | | | |
| Baseline CNN | 55.7± 0.4 | 57.8± 0.4 | 36.6± 0.4 | 41.0± 0.4 | 25.5± 0.4 | √ |
| DNABERT2 (SOTA) | 77.0± 0.1 | 75.9± 0.1 | 65.9± 0.1 | 75.8± 0.1 | 50.4± 0.1 | × |
| HyenaDNA | 66.7± 0.4 | 68.5± 0.4 | 57.4± 0.4 | 50.2± 0.4 | 38.4± 0.4 | × |
| Baseline LegNet | 60.2± 0.5 | 62.0± 0.5 | 38.5± 0.5 | 44.2± 0.5 | 33.9± 0.8 | √ |
| LKAN | 61.8± 0.6 | 63.5± 0.6 | 40.0± 0.6 | 45.8± 0.6 | 38.1± 0.8 | √ |
| CKAN | 60.8± 0.7 | 62.8± 0.7 | 39.5± 0.7 | 44.8± 0.7 | 36.2± 0.9 | √ |

**Table S8.** Classification results for GUE *(Continued from previous page)*.

| | | | | | | |
|---|---|---|---|---|---|---|
| | | | **PROMOTER DETECTION** | | | |
| | | | *ALL* | | | |
| BASELINE CNN | 84.4± 0.5 | 86.0± 0.5 | 77.8± 0.5 | 75.6± 0.5 | 75.7± 0.5 | √ |
| DNABERT2 (SOTA) | 96.0± 0.5 | 95.0± 0.2 | 90.4± 0.2 | 88.6± 0.2 | 86.8± 0.2 | × |
| HYENADNA | 80.3± 0.2 | 65.4± 0.2 | 45.4± 0.2 | 53.7± 0.2 | 47.5± 0.5 | √ |
| BASELINE LEGNET | 85.2± 0.7 | 85.5± 0.7 | 78.2± 0.7 | 76.0± 0.7 | 75.9± 1.1 | √ |
| LKAN | 88.0± 0.8 | 87.8± 0.8 | 81.5± 0.8 | 79.2± 0.8 | 81.2± 1.4 | √ |
| CKAN | 86.5± 0.9 | 86.8± 0.9 | 79.1± 0.9 | 77.1± 0.9 | 77.2± 1.2 | √ |
| | | | *NOTATA* | | | |
| BASELINE CNN | 88.3± 0.5 | 88.2± 0.4 | 80.8± 0.4 | 88.6± 0.4 | 85.1± 0.5 | √ |
| DNABERT2 (SOTA) | 96.0± 0.2 | 95.2± 0.3 | 90.4± 0.2 | 98.6± 0.2 | 94.3± 0.3 | × |
| HYENADNA | 82.0± 0.4 | 81.5± 0.4 | 45.0± 0.4 | 73.0± 0.4 | 52.2± 0.5 | √ |
| BASELINE LEGNET | 84.2± 0.5 | 82.0± 0.4 | 79.0± 0.5 | 87.3± 0.4 | 88.2± 1.1 | √ |
| LKAN | 91.2± 0.6 | 90.3± 0.6 | 87.0± 0.5 | 88.0± 0.6 | 89.9± 0.7 | √ |
| CKAN | 85.5± 0.5 | 84.2± 0.5 | 86.3± 0.5 | 89.4± 0.6 | 87.1± 1.2 | √ |
| | | | *TATA* | | | |
| BASELINE CNN | 70.1± 0.5 | 71.3± 0.5 | 68.7± 0.6 | 69.5± 0.4 | 70.3± 0.5 | √ |
| DNABERT2 (SOTA) | 91.2± 0.2 | 92.5± 0.2 | 89.1± 0.2 | 90.3± 0.1 | 71.6± 0.2 | × |
| HYENADNA | 55.4± 0.1 | 86.2± 0.1 | 32.5± 0.1 | 13.9± 0.1 | 5.34± 0.1 | √ |
| BASELINE LEGNET | 77.3± 0.5 | 78.1± 0.4 | 74.2± 0.5 | 75.5± 0.3 | 56.1± 0.5 | √ |
| LKAN | 80.2± 0.4 | 81.5± 0.4 | 78.0± 0.4 | 79.1± 0.3 | 58.2± 0.8 | √ |
| CKAN | 78.9± 0.5 | 80.0± 0.4 | 76.5± 0.4 | 77.4± 0.3 | 57.9± 0.9 | √ |

**Table S9.** Classification results for GUE *(Continued from previous page)*.

| | | | | | | |
|---|---|---|---|---|---|---|
| | | | **Core Promoter Detection** | | | |
| | | | *ALL* | | | |
| Baseline CNN | 68.1± 0.4 | 69.3± 0.5 | 65.4± 0.5 | 66.3± 0.4 | 58.1± 0.4 | √ |
| DNABERT2 (SOTA) | 80.2± 0.6 | 81.8± 0.6 | 77.5± 0.6 | 78.8± 0.6 | 69.5± 0.6 | × |
| HyenaDNA | 73.5± 0.5 | 74.0± 0.5 | 40.2± 0.5 | 21.0± 0.5 | 37.0± 0.5 | √ |
| Baseline LegNet | 72.0± 0.8 | 73.2± 0.7 | 69.1± 0.6 | 70.0± 0.6 | 64.8± 0.9 | √ |
| LKAN | 76.1± 0.5 | 77.4± 0.5 | 73.6± 0.5 | 74.2± 0.4 | 68.2± 0.6 | √ |
| CKAN | 74.8± 0.7 | 75.9± 0.6 | 71.9± 0.7 | 72.8± 0.5 | 66.2± 1.1 | √ |
| | | | *NOTATA* | | | |
| Baseline CNN | 61.3± 0.2 | 62.0± 0.2 | 58.9± 0.2 | 59.5± 0.2 | 60.0± 0.2 | √ |
| DNABERT2 (SOTA) | 78.5± 0.2 | 79.1± 0.2 | 75.3± 0.2 | 76.4± 0.2 | 68.0± 0.2 | × |
| HyenaDNA | 70.4± 0.2 | 71.0± 0.2 | 27.1± 0.2 | 48.0± 0.2 | 35.4± 0.2 | √ |
| Baseline LegNet | 73.5± 0.6 | 74.0± 0.5 | 70.5± 0.6 | 71.0± 0.4 | 64.5± 0.7 | √ |
| LKAN | 76.3± 0.5 | 77.1± 0.5 | 73.6± 0.5 | 74.3± 0.4 | 67.2± 0.8 | √ |
| CKAN | 74.2± 0.6 | 75.0± 0.6 | 71.2± 0.6 | 72.0± 0.5 | 63.1± 1.1 | √ |
| | | | *TATA* | | | |
| Baseline CNN | 69.6± 0.2 | 70.2± 0.2 | 66.5± 0.2 | 67.0± 0.2 | 69.3± 0.2 | √ |
| DNABERT2 (SOTA) | 81.3± 0.2 | 82.1± 0.2 | 78.0± 0.2 | 79.1± 0.2 | 74.2± 0.2 | × |
| HyenaDNA | 77.1± 0.2 | 78.0± 0.2 | 73.2± 0.2 | 74.0± 0.2 | 72.9± 0.2 | √ |
| Baseline LegNet | 75.4± 0.7 | 76.0± 0.6 | 72.5± 0.6 | 73.2± 0.5 | 72.3± 0.9 | √ |
| LKAN | 80.5± 0.7 | 81.2± 0.7 | 77.3± 0.7 | 78.0± 0.5 | 78.2± 0.9 | √ |
| CKAN | 78.4± 0.8 | 79.0± 0.7 | 74.9± 0.8 | 75.5± 0.6 | 74.7± 1.2 | √ |

**Table S10.** Classification results for GUE *(Continued from previous page)* .

| Transcription Factor Prediction (Human) | | | | | | |
|---|---|---|---|---|---|---|
| *0* | | | | | | |
| Model | Acc | AUC | PR | F1 | MCC | Better? |
| Baseline CNN | 55.5± 0.2 | 56.0± 0.2 | 53.0± 0.2 | 53.5± 0.2 | 54.0± 0.2 | √ |
| DNABERT2 (SOTA) | 87.2± 0.2 | 88.1± 0.2 | 83.0± 0.2 | 84.5± 0.2 | 84.4± 0.2 | × |
| HyenaDNA | 75.0± 0.3 | 76.0± 0.3 | 60.8± 0.4 | 61.2± 0.3 | 62.3± 0.3 | √ |
| Baseline LegNet | 76.5± 0.6 | 77.0± 0.5 | 70.5± 0.6 | 71.0± 0.5 | 65.1± 0.7 | √ |
| LKAN | 80.0± 0.5 | 81.2± 0.5 | 75.4± 0.5 | 76.0± 0.4 | 67.4± 0.6 | √ |
| CKAN | 78.1± 0.7 | 78.9± 0.6 | 72.3± 0.7 | 73.1± 0.5 | 65.5± 0.9 | √ |
| *1* | | | | | | |
| Baseline CNN | 64.1± 0.2 | 64.7± 0.2 | 61.3± 0.2 | 62.0± 0.2 | 63.2± 0.2 | √ |
| DNABERT2 (SOTA) | 88.5± 0.2 | 89.1± 0.2 | 85.4± 0.2 | 86.0± 0.2 | 87.2± 0.2 | × |
| HyenaDNA | 77.2± 0.2 | 78.0± 0.2 | 66.5± 0.2 | 67.1± 0.2 | 67.8± 0.2 | √ |
| Baseline LegNet | 77.5± 1.0 | 78.0± 1.1 | 71.0± 1.0 | 71.5± 1.1 | 69.9± 1.0 | √ |
| LKAN | 81.5± 0.8 | 82.2± 0.7 | 77.0± 0.7 | 77.5± 0.6 | 72.4± 0.8 | √ |
| CKAN | 78.0± 1.2 | 78.5± 1.0 | 73.2± 1.1 | 73.5± 0.9 | 66.3± 1.3 | √ |
| *2* | | | | | | |
| Baseline CNN | 44.5± 0.2 | 45.1± 0.2 | 42.5± 0.2 | 43.0± 0.2 | 45.2± 0.2 | √ |
| DNABERT2 (SOTA) | 83.6± 0.2 | 84.2± 0.2 | 80.8± 0.2 | 81.0± 0.2 | 84.0± 0.2 | × |
| HyenaDNA | 46.2± 0.2 | 46.6± 0.2 | 43.1± 0.2 | 43.5± 0.2 | 47.0± 0.2 | √ |
| Baseline LegNet | 48.2± 0.3 | 48.8± 0.3 | 45.0± 0.4 | 45.5± 0.4 | 49.6± 0.4 | √ |
| LKAN | 52.3± 0.6 | 53.1± 0.5 | 49.8± 0.5 | 50.0± 0.5 | 54.3± 0.7 | √ |
| CKAN | 46.5± 0.6 | 47.0± 0.6 | 43.7± 0.7 | 44.0± 0.6 | 47.2± 0.7 | √ |
| *3* | | | | | | |
| Baseline CNN | 28.0± 0.2 | 28.5± 0.2 | 26.2± 0.2 | 26.5± 0.2 | 29.8± 0.2 | √ |
| DNABERT2 (SOTA) | 78.0± 0.2 | 79.0± 0.2 | 77.5± 0.2 | 77.8± 0.2 | 79.9± 0.2 | × |
| HyenaDNA | 40.0± 0.2 | 40.5± 0.2 | 38.0± 0.2 | 38.2± 0.2 | 41.8± 0.2 | √ |
| Baseline LegNet | 42.0± 0.4 | 42.5± 0.4 | 41.0± 0.4 | 41.2± 0.4 | 43.2± 0.5 | √ |
| LKAN | 48.2± 0.5 | 48.8± 0.5 | 48.0± 0.5 | 45.3± 0.5 | 46.8± 0.5 | √ |
| CKAN | 43.0± 0.8 | 43.5± 0.8 | 42.0± 0.9 | 42.2± 0.8 | 44.4± 0.9 | √ |
| *4* | | | | | | |
| Baseline CNN | 70.1± 0.2 | 69.8± 0.2 | 69.6± 0.2 | 70.0± 0.2 | 61.5± 0.2 | √ |
| DNABERT2 (SOTA) | 87.9± 0.2 | 88.1± 0.2 | 88.0± 0.2 | 87.8± 0.2 | 87.8± 0.2 | × |
| HyenaDNA | 79.5± 0.2 | 80.2± 0.2 | 78.9± 0.2 | 80.1± 0.2 | 61.2± 0.2 | × |
| Baseline LegNet | 72.6± 0.8 | 73.2± 0.8 | 72.5± 0.8 | 72.7± 0.8 | 72.6± 0.8 | √ |
| LKAN | 73.4± 0.5 | 73.0± 0.5 | 73.5± 0.5 | 73.6± 0.5 | 72.9± 0.5 | √ |
| CKAN | 71.2± 0.9 | 71.5± 0.9 | 71.0± 0.9 | 71.4± 0.9 | 71.3± 0.9 | √ |

**Table S11.** Classification results for GUE *(Continued from previous page)* .

| Virus | | | | | | |
|---|---|---|---|---|---|---|
| Model | Acc | AUC | PR | F1 | MCC | Better? |
| Baseline CNN | 65.2± 0.2 | 64.7± 0.2 | 64.5± 0.2 | 64.8± 0.2 | 22.2± 0.2 | √ |
| DNABERT2 (SOTA) | 88.1± 0.4 | 88.4± 0.4 | 88.0± 0.4 | 88.2± 0.4 | 71.0± 0.4 | × |
| HyenaDNA | 74.8± 0.2 | 75.2± 0.2 | 74.5± 0.2 | 74.9± 0.2 | 23.3± 0.2 | √ |
| Baseline LegNet | 70.3± 0.9 | 70.1± 0.9 | 70.2± 0.9 | 70.0± 0.9 | 50.3± 0.9 | √ |
| LKAN | 75.5± 1.5 | 75.0± 1.5 | 75.2± 1.5 | 75.3± 1.5 | 58.2± 1.5 | √ |
| CKAN | 71.4± 0.6 | 71.2± 0.6 | 71.1± 0.6 | 71.3± 0.6 | 50.5± 0.6 | √ |
| Splice | | | | | | |
| Baseline CNN | 77.7± 0.2 | 78.1± 0.2 | 77.5± 0.2 | 77.6± 0.2 | 77.7± 0.2 | √ |
| DNABERT2 (SOTA) | 85.0± 0.2 | 85.2± 0.2 | 84.9± 0.2 | 85.1± 0.2 | 85.0± 0.2 | × |
| HyenaDNA | 72.7± 0.2 | 73.0± 0.2 | 72.5± 0.2 | 72.8± 0.2 | 72.7± 0.2 | √ |
| Baseline LegNet | 78.2± 0.8 | 78.4± 0.8 | 77.9± 0.8 | 78.1± 0.8 | 78.2± 0.8 | √ |
| LKAN | 80.0± 0.8 | 80.1± 0.8 | 79.8± 0.8 | 79.9± 0.8 | 80.0± 0.8 | √ |
| CKAN | 75.2± 0.7 | 75.5± 0.7 | 75.0± 0.7 | 75.1± 0.7 | 75.2± 0.7 | √ |

**Table S12.** Classification results for GUE *(Continued from previous page)*.

| Transcription Factor Prediction (Mouse) | | | | | | |
|---|---|---|---|---|---|---|
| *0* | | | | | | |
| Model | Acc | AUC | PR | F1 | MCC | Better? |
| Baseline CNN | 51.1± 0.2 | 50.8± 0.2 | 51.0± 0.2 | 50.9± 0.2 | 31.1± 0.2 | √ |
| DNABERT2 (SOTA) | 81.6± 0.2 | 81.9± 0.2 | 81.5± 0.2 | 81.7± 0.2 | 81.6± 0.2 | × |
| HyenaDNA | 55.6± 0.2 | 65.8± 0.2 | 55.5± 0.2 | 45.7± 0.2 | 35.6± 0.2 | √ |
| Baseline LegNet | 58.4± 0.7 | 58.2± 0.7 | 48.3± 0.7 | 58.2± 0.7 | 38.2± 0.7 | √ |
| LKAN | 70.2± 0.6 | 67.0± 0.6 | 50.1± 0.6 | 60.0± 0.6 | 40.1± 0.6 | √ |
| CKAN | 39.1± 0.9 | 39.0± 0.9 | 39.2± 0.9 | 39.0± 0.9 | 39.0± 0.9 | √ |
| *1* | | | | | | |
| Baseline CNN | 69.7± 0.2 | 60.0± 0.2 | 59.5± 0.2 | 59.6± 0.2 | 59.7± 0.2 | √ |
| DNABERT2 (SOTA) | 92.6± 0.2 | 92.8± 0.2 | 92.5± 0.2 | 92.7± 0.2 | 92.6± 0.2 | × |
| HyenaDNA | 89.5± 0.2 | 84.8± 0.2 | 80.3± 0.2 | 80.6± 0.2 | 80.5± 0.2 | √ |
| Baseline LegNet | 81.0± 0.7 | 81.2± 0.7 | 80.9± 0.7 | 81.1± 0.7 | 81.0± 0.7 | √ |
| LKAN | 82.8± 0.6 | 83.0± 0.6 | 82.7± 0.6 | 82.8± 0.6 | 82.8± 0.6 | √ |
| CKAN | 80.1± 0.8 | 80.3± 0.8 | 79.9± 0.8 | 80.0± 0.8 | 80.1± 0.8 | √ |
| *2* | | | | | | |
| Baseline CNN | 63.1± 0.2 | 63.3± 0.2 | 63.0± 0.2 | 63.2± 0.2 | 63.1± 0.2 | √ |
| DNABERT2 (SOTA) | 92.3± 0.2 | 92.5± 0.2 | 92.2± 0.2 | 92.4± 0.2 | 92.3± 0.2 | × |
| HyenaDNA | 65.3± 0.2 | 65.5± 0.2 | 65.2± 0.2 | 65.4± 0.2 | 65.3± 0.2 | √ |
| Baseline LegNet | 79.8± 0.6 | 80.0± 0.6 | 79.7± 0.6 | 79.9± 0.6 | 79.8± 0.6 | √ |
| LKAN | 85.5± 0.8 | 85.7± 0.8 | 85.4± 0.8 | 85.6± 0.8 | 85.5± 0.8 | √ |
| CKAN | 81.2± 0.7 | 81.4± 0.7 | 81.1± 0.7 | 81.3± 0.7 | 81.2± 0.7 | √ |
| *3* | | | | | | |
| Baseline CNN | 55.4± 0.2 | 50.5± 0.2 | 45.3± 0.2 | 55.4± 0.2 | 45.4± 0.2 | √ |
| DNABERT2 (SOTA) | 84.9± 0.2 | 85.0± 0.2 | 84.8± 0.2 | 84.9± 0.2 | 84.9± 0.2 | × |
| HyenaDNA | 64.2± 0.2 | 62.3± 0.2 | 64.1± 0.2 | 54.2± 0.2 | 54.2± 0.2 | √ |
| Baseline LegNet | 77.6± 1.2 | 77.8± 1.2 | 77.5± 1.2 | 77.7± 1.2 | 77.6± 1.2 | √ |
| LKAN | 81.4± 0.7 | 78.5± 0.7 | 79.3± 0.7 | 80.4± 0.7 | 81.4± 0.7 | √ |
| CKAN | 77.2± 0.9 | 79.0± 0.9 | 79.0± 0.9 | 79.2± 0.9 | 79.2± 0.9 | √ |
| *4* | | | | | | |
| Baseline CNN | 47.1± 0.2 | 49.2± 0.2 | 27.0± 0.2 | 36.1± 0.2 | 27.1± 0.2 | √ |
| DNABERT2 (SOTA) | 76.1± 0.2 | 76.2± 0.2 | 76.0± 0.2 | 76.1± 0.2 | 76.1± 0.2 | × |
| HyenaDNA | 49.1± 0.2 | 51.2± 0.2 | 19.0± 0.2 | 24.1± 0.2 | 19.1± 0.2 | √ |
| Baseline LegNet | 42.6± 0.7 | 42.7± 0.7 | 42.5± 0.7 | 42.6± 0.7 | 42.6± 0.7 | √ |
| LKAN | 48.7± 0.5 | 48.8± 0.5 | 48.6± 0.5 | 48.7± 0.5 | 48.7± 0.5 | √ |
| CKAN | 40.1± 0.7 | 40.2± 0.7 | 40.0± 0.7 | 40.1± 0.7 | 40.1± 0.7 | √ |

**Table S13.** Classification results for GUE+

| Epigenetic Marks Prediction | | | | | | |
|---|---|---|---|---|---|---|
| *GM12878* | | | | | | |
| Model | Acc | AUC | PR | F1 | MCC | Better? |
| Baseline CNN | 57.1± 0.2 | 57.2± 0.2 | 32.0± 0.2 | 30.1± 0.2 | 27.1± 0.2 | √ |
| DNABERT2 (SOTA) | 76.1± 0.2 | 76.2± 0.2 | 76.0± 0.2 | 76.1± 0.2 | 76.2± 0.2 | × |
| HyenaDNA | 61.8± 0.2 | 61.9± 0.2 | 61.7± 0.2 | 61.8± 0.2 | 61.9± 0.2 | × |
| Baseline LegNet | 56.5± 1.1 | 56.7± 1.1 | 36.4± 1.1 | 36.6± 1.1 | 36.6± 1.1 | √ |
| LKAN | 68.0± 0.9 | 71.1± 0.9 | 40.9± 0.9 | 51.0± 0.9 | 41.2± 0.9 | √ |
| CKAN | 56.8± 1.2 | 57.0± 1.2 | 36.6± 1.2 | 37.0± 1.2 | 37.0± 1.2 | √ |
| *HELA-S3* | | | | | | |
| Baseline CNN | 68.1± 0.2 | 60.2± 0.2 | 51.3± 0.2 | 49.0± 0.2 | 28.7± 0.2 | √ |
| DNABERT2 (SOTA) | 92.3± 0.2 | 93.2± 0.2 | 92.5± 0.2 | 92.0± 0.2 | 79.2± 0.2 | × |
| HyenaDNA | 82.5± 0.2 | 83.1± 0.2 | 82.7± 0.2 | 82.3± 0.2 | 72.2± 0.2 | × |
| Baseline LegNet | 77.8± 1.0 | 78.5± 1.0 | 77.9± 1.0 | 68.0± 1.0 | 34.3± 1.0 | √ |
| LKAN | 84.2± 1.2 | 85.0± 1.2 | 74.5± 1.2 | 84.8± 1.2 | 40.8± 1.2 | √ |
| CKAN | 72.3± 0.9 | 73.1± 0.9 | 62.7± 0.9 | 62.8± 0.9 | 31.3± 0.9 | √ |
| *HUVEC* | | | | | | |
| Baseline CNN | 59.2± 0.2 | 51.0± 0.2 | 50.5± 0.2 | 49.5± 0.2 | 28.1± 0.2 | √ |
| DNABERT2 (SOTA) | 92.1± 0.2 | 93.5± 0.2 | 92.8± 0.2 | 92.3± 0.2 | 83.5± 0.2 | × |
| HyenaDNA | 81.3± 0.2 | 82.7± 0.2 | 82.1± 0.2 | 81.8± 0.2 | 73.1± 0.2 | × |
| Baseline LegNet | 68.7± 0.7 | 69.2± 0.7 | 68.5± 0.7 | 68.0± 0.7 | 37.2± 0.7 | √ |
| LKAN | 72.3± 0.9 | 73.1± 0.9 | 72.5± 0.9 | 72.8± 0.9 | 39.6± 0.9 | √ |
| CKAN | 62.9± 0.8 | 63.5± 0.8 | 63.0± 0.8 | 63.2± 0.8 | 30.2± 0.8 | √ |
| *IMR90* | | | | | | |
| Baseline CNN | 57.1± 0.2 | 58.0± 0.2 | 47.5± 0.2 | 46.8± 0.2 | 27.2± 0.2 | √ |
| DNABERT2 (SOTA) | 91.2± 0.2 | 92.5± 0.2 | 92.1± 0.2 | 91.8± 0.2 | 86.7± 0.2 | × |
| HyenaDNA | 83.1± 0.2 | 84.0± 0.2 | 83.5± 0.2 | 83.2± 0.2 | 79.5± 0.2 | × |
| Baseline LegNet | 67.8± 0.9 | 68.4± 0.9 | 67.9± 0.9 | 67.5± 0.9 | 34.5± 0.9 | √ |
| LKAN | 78.3± 0.9 | 80.0± 0.9 | 71.7± 0.9 | 71.5± 0.9 | 42.0± 0.9 | √ |
| CKAN | 60.2± 1.3 | 61.0± 1.3 | 60.8± 1.3 | 60.5± 1.3 | 33.3± 1.3 | √ |
| *K692* | | | | | | |
| Baseline CNN | 69.5± 0.2 | 70.3± 0.2 | 49.9± 0.2 | 49.4± 0.2 | 27.1± 0.2 | √ |
| DNABERT2 (SOTA) | 93.1± 0.2 | 93.8± 0.2 | 93.4± 0.2 | 93.0± 0.2 | 92.9± 0.2 | × |
| HyenaDNA | 87.2± 0.2 | 88.0± 0.2 | 87.6± 0.2 | 87.1± 0.2 | 86.4± 0.2 | × |
| Baseline LegNet | 78.9± 0.9 | 79.5± 0.9 | 69.0± 0.9 | 68.7± 0.9 | 38.2± 0.9 | √ |
| LKAN | 83.4± 0.7 | 84.0± 0.7 | 73.7± 0.7 | 83.5± 0.7 | 44.4± 0.7 | √ |
| CKAN | 70.1± 1.1 | 70.8± 1.1 | 60.5± 1.1 | 60.0± 1.1 | 37.9± 1.1 | √ |
| *NHEK* | | | | | | |
| Baseline CNN | 68.5± 0.2 | 69.0± 0.2 | 48.7± 0.2 | 48.3± 0.2 | 27.4± 0.2 | √ |
| DNABERT2 (SOTA) | 91.2± 0.2 | 91.8± 0.2 | 91.5± 0.2 | 91.1± 0.2 | 73.7± 0.2 | × |
| HyenaDNA | 84.3± 0.2 | 85.0± 0.2 | 84.7± 0.2 | 84.2± 0.2 | 68.6± 0.2 | × |
| Baseline LegNet | 77.1± 1.1 | 77.7± 1.1 | 57.3± 1.1 | 67.0± 1.1 | 33.7± 1.1 | √ |
| LKAN | 82.4± 0.9 | 83.1± 0.9 | 62.8± 0.9 | 72.3± 0.9 | 40.1± 0.9 | √ |
| CKAN | 75.7± 0.2 | 76.3± 0.2 | 56.0± 0.2 | 75.6± 0.2 | 36.7± 0.2 | √ |

**Table S14.** Classification results for GUE+

| Epigenetic Marks Prediction | | | | | | |
|---|---|---|---|---|---|---|
| Model | Acc | AUC | PR | F1 | MCC | Better? |
| | | | *FUNGI* | | | |
| Baseline CNN | 54.4± 0.2 | 54.6± 0.2 | 44.3± 0.2 | 44.5± 0.2 | 44.5± 0.2 | √ |
| DNABERT2 (SOTA) | 68.4± 0.2 | 68.6± 0.2 | 48.3± 0.2 | 58.5± 0.2 | 48.5± 0.2 | × |
| HyenaDNA | 55.6± 0.2 | 55.8± 0.2 | 45.5± 0.2 | 45.7± 0.2 | 45.7± 0.2 | × |
| Baseline LegNet | 51.0± 0.6 | 51.2± 0.6 | 50.8± 0.6 | 51.1± 0.6 | 41.1± 0.6 | √ |
| LKAN | 55.4± 1.0 | 55.6± 1.0 | 55.3± 1.0 | 55.5± 1.0 | 45.5± 1.0 | √ |
| CKAN | 50.3± 1.2 | 50.5± 1.2 | 50.2± 1.2 | 50.4± 1.2 | 40.4± 1.2 | √ |
| | | | *VIRUS* | | | |
| Baseline CNN | 49.4± 0.2 | 50.4± 0.2 | 27.4± 0.2 | 33.3± 0.2 | 29.2± 0.2 | √ |
| DNABERT2 (SOTA) | 98.4± 0.2 | 96.6± 0.2 | 94.5± 0.2 | 94.9± 0.2 | 93.0± 0.2 | × |
| HyenaDNA | 84.1± 0.2 | 80.9± 0.2 | 67.8± 0.2 | 80.5± 0.2 | 72.9± 0.2 | × |
| Baseline LegNet | 70.5± 0.8 | 71.5± 0.7 | 40.8± 0.9 | 49.6± 0.6 | 20.7± 0.5 | √ |
| LKAN | 80.3± 1.2 | 82.4± 1.0 | 55.0± 1.0 | 67.3± 1.0 | 35.3± 1.9 | √ |
| CKAN | 71.1± 1.5 | 73.6± 1.5 | 45.6± 1.5 | 51.5± 1.6 | 22.1± 1.8 | √ |

**Table S15.** Evaluating the impact of Grid size parameters (GS) in Linear Kolmogorov Arnold Network layers along with number of blocks with replaced linear layer. $N$ denote a number of blocks incorporated with LKANs.

| Model | GS=3 | GS=5 | GS=7 | GS=10 |
|---|---|---|---|---|
| LKAN (N=1) | 66.1 | 67.5 | 69.0 | 70.5 |
| LKAN (N=2) | 67.0 | 68.5 | 70.0 | 71.4 |
| LKAN (N=3) | 67.9 | 70.8 | 71.5 | 72.5 |
| LKAN (N=4) | 68.0 | 71.2 | 72.0 | 72.7 |
| LKAN (N=5) | 70.3 | 71.0 | 72.7 | 73.0 |
| LKAN (N=6) | 71.0 | 72.6 | 73.1 | 73.5 |
| CKAN (N=1) | 63.8 | 64.5 | 65.2 | 65.8 |
| CKAN (N=2) | 64.1 | 65.3 | 66.3 | 66.5 |
| CKAN (N=3) | 65.0 | 66.1 | 66.6 | 67.4 |
| CKAN (N=4) | 65.8 | 66.5 | 67.7 | 68.0 |
| CKAN (N=5) | 66.5 | 67.2 | 68.1 | 68.5 |
| CKAN (N=6) | 67.6 | 68.0 | 69.8 | 70.9 |

**Table S16.** Evaluating the time scaling of number of blocks with replaced linear layer. $N$ denote a number of blocks incorporated with LKANs and CKANs respectively.

| Model | Time per epoch (min) |
|---|---|
| LKAN (N=1) | 4.0 |
| LKAN (N=2) | 5.7 |
| LKAN (N=3) | 5.8 |
| LKAN (N=4) | 7.0 |
| LKAN (N=5) | 9.1 |
| LKAN (N=6) | 12.4 |
| CKAN (N=1) | 26.2 |
| CKAN (N=2) | 30.1 |
| CKAN (N=3) | 34.5 |
| CKAN (N=4) | 44.6 |
| CKAN (N=5) | 51.0 |
| CKAN (N=6) | 55.7 |
